# Sampling of Structure and Sequence Space of Small Protein Folds

**DOI:** 10.1101/2021.03.10.434454

**Authors:** T Linsky, K Noble, A Tobin, R Crow, Lauren Carter, J Urbauer, D Baker, EM Strauch

## Abstract

Nature only samples a small fraction in sequence space, yet many more amino acid combinations can fold into stable proteins. Furthermore, small structural variations in a single fold, which may only be a few amino acids different from the next homolog, define their molecular function. Hence, to design proteins with novel molecular functionalities, such as molecular recognition, methods to control and sample shape diversity are necessary. To explore this space, we developed and experimentally validated a computational platform that can design a wide variety of small protein folds while sampling high shape diversity. We designed and evaluated about 30,000 de novo protein designs of 7 different folds. Among these designs, about 6,200 stable proteins were identified, with predicted structures having first-of-its-kind minimalized thioredoxin. Obtained data revealed more protein folding rules, such as helix connecting loops, which were in nature. Beyond providing a resource database for protein engineering, our data presents a large training data set for machine learning. We developed a high-accuracy classifier to predict the stability of our designed proteins. The methods and the wide range of new protein shapes provide a basis for the design of new protein function without compromising stability.

## Main

Proteins are part of most biological processes and function as catalysts, messengers and transporters to name only a few of their tasks. Their sequences determine their structures and thereby also their molecular role. Yet, natural sequence space only covers a small fraction of possible proteins^1^. Furthermore, evolution of molecular function generally occurs through the diversification of a rather small number of known protein families, highlighting the power of shape diversity to increase functional diversity^2^. Hence, the sampling and control of small structural variations with high accuracy represents an important advancement in our ability to design new proteins with new functionalities. So far, a few *de novo* designed globular folds^3-5^ have been generated, but not much structural diversity within a given fold has been purposely sampled and importantly, experimentally verified, largely due to the lack of a versatile computational infrastructure. Recent advances demonstrated that the exploration of a Loop-Helix-Loop motif enables further sampling of existing folds^6^; however, here we go beyond and describe how to exhaust the plasticity of small protein folds at large scale, sampling each secondary structure and loop connector to provide diversity within the given restraints (Fig.1, S1).

With the advent of oligonucleotide synthesis technology^7^, thousands of short genes (currently encoding up to 64 amino acids, but sizes are increasing as the technology is being further developed) can be affordably manufactured as single oligonucleotides in large pools; proteins up to 110 amino acids can also be synthesized via assembly methods^8^. Combined with yeast surface display methods, this has enabled the rapid, high-throughput experimental evaluation of the stability of thousands of proteins at a time^1^. However, computational methods for *de novo* design of monomeric proteins in Rosetta has so far been limited to a relatively small number of helical bundle and simple beta-strand-containing topologies. This limitation is driven by a reliance on static, pre-determined ranges of secondary structure lengths and backbone torsion angles required to form the desired secondary and tertiary structural elements of the protein. The 5 distinct areas of torsion angle distributions found in natural proteins as illustrated in the Ramachandran plot (Fig. 1C) can be described by a five-letter alphabet: ABEG0^9^ (Fig. 1C). This extends the secondary structure description beyond the simple alpha-helical (“A”), beta-sheet (“B”) and loop regions to more precisely describe the loop conformational space. Design protocols up to this point have been limited to using fixed and predefined ABEG0 sequences (as a “blueprint”) for a design trajectory. Proteins are then folded *in silico* by inserting structured fragments curated from the PDB^3, 10, 11^ into an extended chain. However, this approach results in costly computation as it tries to fold an entire protein either at once or in two steps that were manually predetermined by an expert. This limits both the scalability of the approach for larger proteins, and places restrictions on the sampling of the shape space for a given fold; variations within any of the secondary structure lengths are not possible unless a new blueprint file is written, and the trajectory is independently restarted. However, the ability to sample and control small geometric variations of a protein is crucial for the engineering of new functionalities; for instance, new binding proteins variations will have to fit into variously shaped pockets.

**Fig 1.**
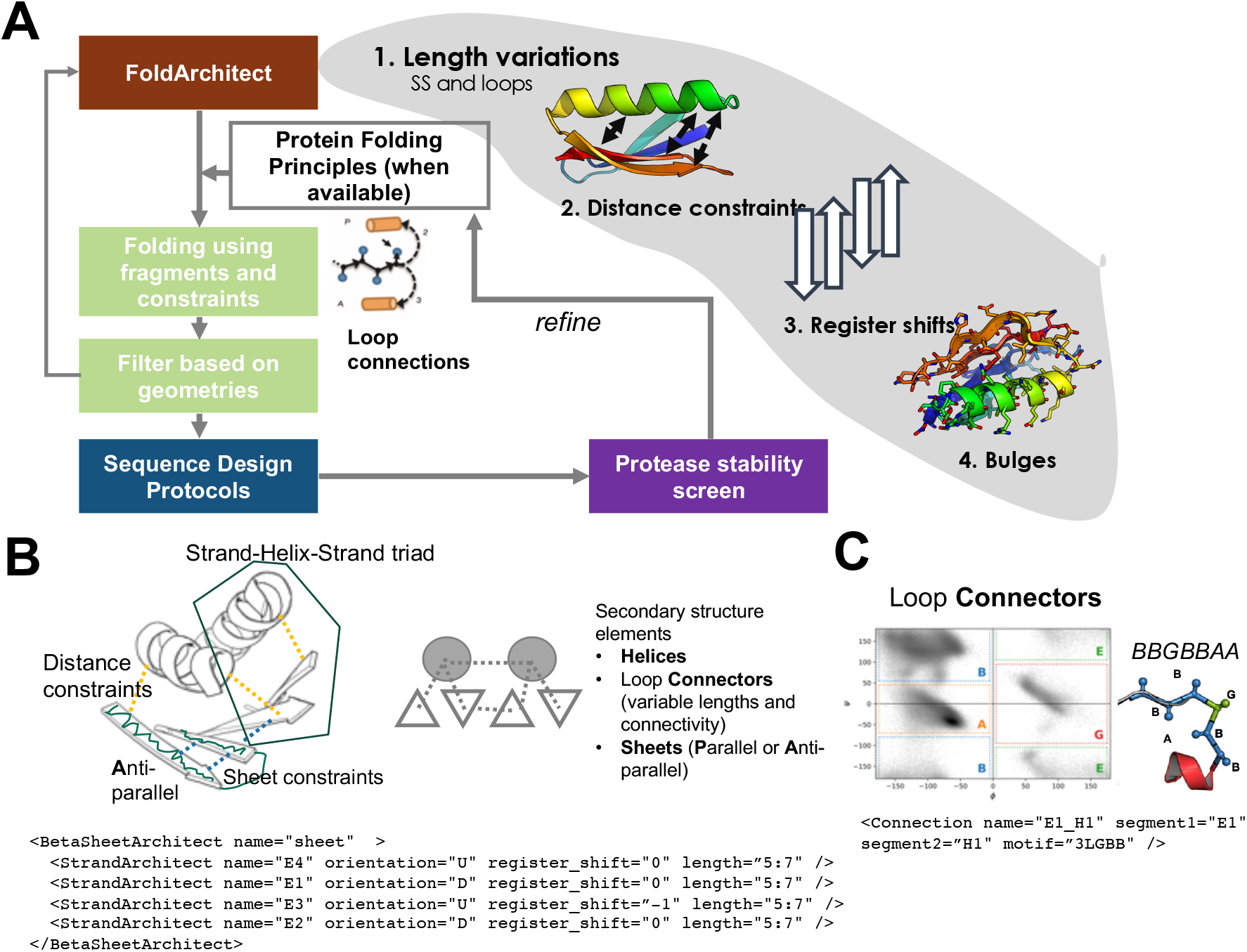
Overview. (**A**) The user can specify for the Fold Architect which ranges of secondary structure (SS) and loop lengths should be sampled, what distance constraints should be met and how it is applied (for instance harmonic constraints), what register shifts should be sampled or whether beta-stands should have a bulge to introduce a larger curvature. We then use the previously reported fragment insertion protocol^1^ and filter for geometrically realistic backbone conformation before designing the sequence of the new poly-valine construct. After sequence design, the decoys are then screened for their stability using varying concentrations of trypsin and chymotrypsin. Information gained from the stable designs was fed back into the FoldArchitect.(**B**) Various distance constraints (yellow lines) and secondary structure element pairing can be encoded in a fold-description file (following an xml format) that enables to record next to length variations also specific or ambiguous distance constraints, pairing of helices or helix-sheet-helix, sheet orientations or bulge insertions into beta-sheets. (**C**) Ramachandran plot and the ABEG0 regions and the format of how loop connectors can be specified.

Here, we provide a computational pipeline for the massively parallel design of proteins from scratch to rapidly explore the shape diversity of protein folds and take advantage of high throughput experimental methods to evaluate it. Our approach enables dynamic conformational and topological sampling of folds without prior knowledge of residue-by-residue features. It enables the design of a diverse representation of a given fold and allows sampling during each trajectory of (1) the lengths of each secondary structure element, (2) the distance(s) between secondary structure elements (which can also be assessed as a distribution), (3) the alignment of *α*-helices and *β*-strands and, if more than two strands are involved, their register shifts and lastly (4) the introduction and placement of bulges to introduce curvature into *β*-strands^5^ (Fig.1). The framework automatically applies previously discovered protein folding rules for loop connections^4^ and their sequence biases and is extendable to new structural features. The RosettaScripts XML scheme allows modification of the design strategy without programming knowledge.

## Results

The number of amino acids that can be encoded within a single oligonucleotide in large pools caps the number of independent secondary structured elements within each protein. For a 230 nt oligo, the maximum length of ~64 amino acids allow up to about 6 independent secondary structure elements per fold, sufficient for several diverse fold sub-families. We designed several alpha/beta (protein are structurally composed of alternating *α*-helices and *β*-sheets in which the beta sheets are mostly parallel to each other), alpha-beta(*α*-helices and *β*-sheets that occur separately), and various non-parametric *α*-helical folds: We built 3- and 4-helical bundles, supercoiled 4-helical bundles, beta-grasps, ferredoxins, thioredoxin and two folds not found in nature (Fig. S2).

Each fold is subdivided into segments, which are folded and validated incrementally. During the folding trajectory of each segment, different features such as secondary structure length and loop types are varied dynamically to find the best set of properties for the new segment in the context of the previously folded segments given demanded constraints. This results in a diverse set of backbones for a given fold (Fig. S3). For each beta-stand containing fold, introduced loose distance constraints (Suppl. XML files, Fig. S4). The algorithm automatically generates the beta-strand pairing between neighboring strands and their directions (parallel *versus* antiparallel) can be specified. It samples the different previously pre-defined loop conformations (specific ABEG0 sequences) according to predefined protein folding rules obtained through the design of ferredoxins^6^ and now also this work. After *in silico* folding of the complete protein, we developed two different sequence design protocols. For the first protocol, we sample rotamers of a select set of amino acids choices biased depending on how much the protein was surface exposed. The protocol starts with low repulsive terms in order to find optimal sidechain interactions and slowly increases itto relieve clashes while keeping the strongest interactions intact. As a second protocol, we took advantage of “pair-motifs”. Pair-motifs are side-chain pairings observed in high resolution crystal structures of natural proteins and have previously successfully guided the design of *de novo* oligomeric assemblies^12^. Before executing the same rotamer-based design approach as described for the first one, we introduce first paired-amino acids. This also tremendously reduced the compute time to re-design the resulting poly-valine backbone. Interestingly, it improved also the rate of successful designs for 3-helical bundles significantly (Fig. S5). Likely, because motifs were selected heavily from larger helical protein-protein interfaces^12^.We selected designs for each topology and sequence design protocol based on a set of filter terms and their computed energies (Methods, SI). Each design has a unique three-dimensional conformation and a unique sequence predicted to be near-optimal for that conformation(Fig. S3).

We experimentally characterized 31,500 sequences reflecting these 7 folds, including about 2,300 randomized sequences as negative controls. The pool was subjected to a titration of trypsin and chymotrypsin as previously reported^1^. However, additional selections and duplicates were also performed to improve accuracy (Methods). We fit EC50 values for over 31,180 sequences (including control sequences) for both proteases for which we then calculated a stability score as previously reported^1^ (Fig.S6). To improve comparability between assays, we added five proteins (Fig. 2C) spanning a wide range of previously measured stability scores using the same protease-based stability evaluation procedure and thereby representing a “stability score ladder” as internal control. This ladder allows adjustment for the activity of each enzyme batch. Digestion of the randomized sequences enabled to determine the stability threshold (Fig. S7).

**Figure 2.**
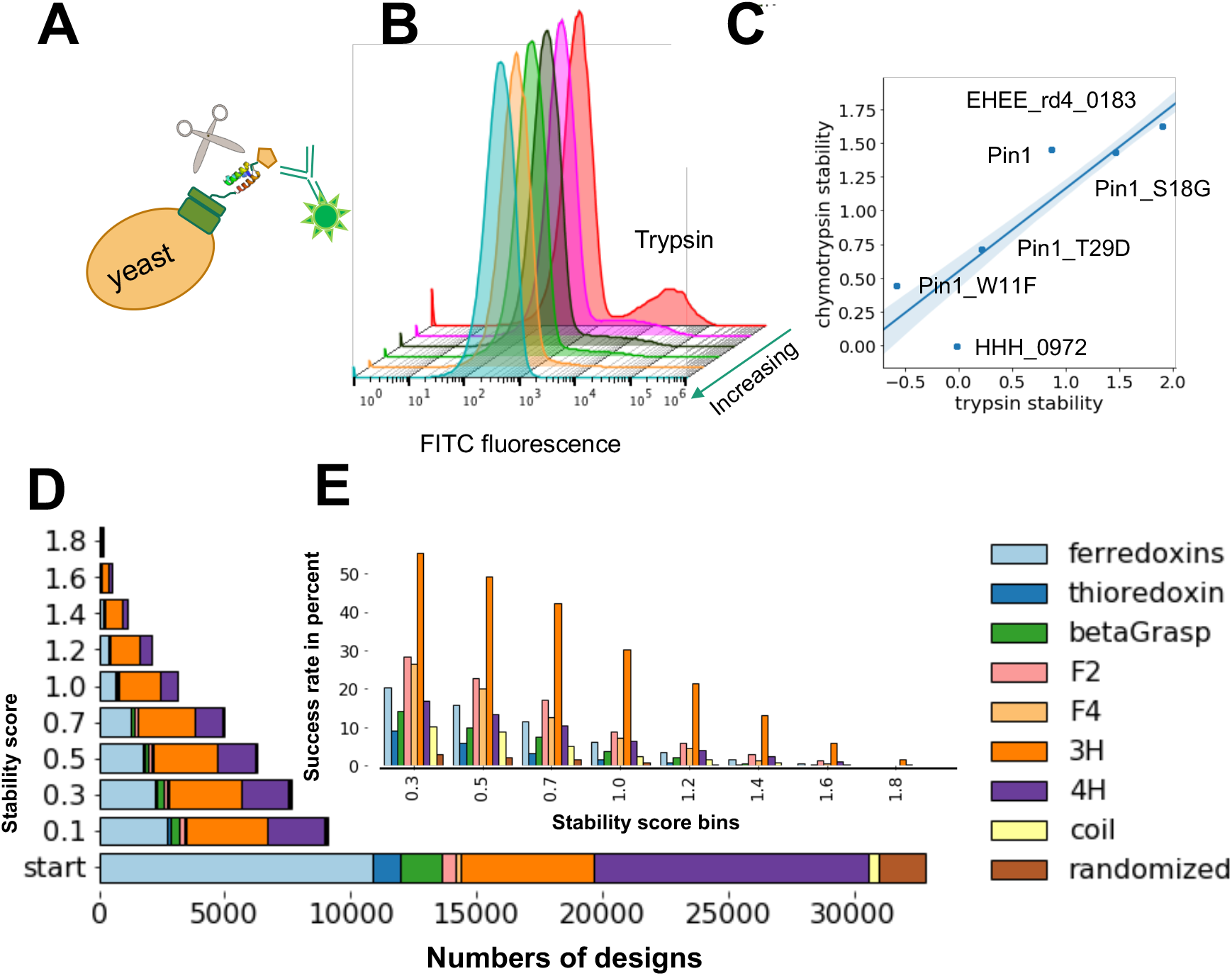
Characterization of designed small proteins using a protease-based thigh-high put screen on the surface of yeast and biochemical analysis of individually expressed proteins. **(A)** Cartoon scheme of yeast surface display experiment. Unfolded proteins are cleaved, and thereby will not be fluorescently labeled. (**B**) Histograms of FITC fluorescence after incubation of yeast cells displaying the designed proteins as a pool with increasing concentration of trypsin; cells were labeled with a c-Myc antibody conjugated to FITC. (**C**) Previously evaluated proteins with known stability scores were included in the pool of the query proteins as a “stability score ladder” to help adjust for protease activities. (**D**) Bar chart of all designs binned at each indicated stability threshold together with (**E**) a bar chart of the success rate of each designed topology for a given stability score bin.

We randomly picked 20 designs with different topologies and stability scores for detailed characterization (Table S1). Of these, 18 proteins expressed soluble in *E. coli*. The F2 and F4 designs appeared to be dimeric as determined by size exclusion chromatography (SEC, Fig. S8). For all other designs we measured the secondary structure of the monomeric protein using circular dichroism (CD) and found them all to be folded. We measured thermostability for several of them (Fig. 3) and found them to be highly stable. We tested one design (ThioFL24) that had a high trypsin stability score, yet a negative score for chymotrypsin. ThioFL24 was soluble and monodisperse (Table S1); however, the CD spectrum indicated that it was only partially folded (Fig. 3), demonstrating that the use of two proteases is important for this high-throughput screen.

**Figure 3.**
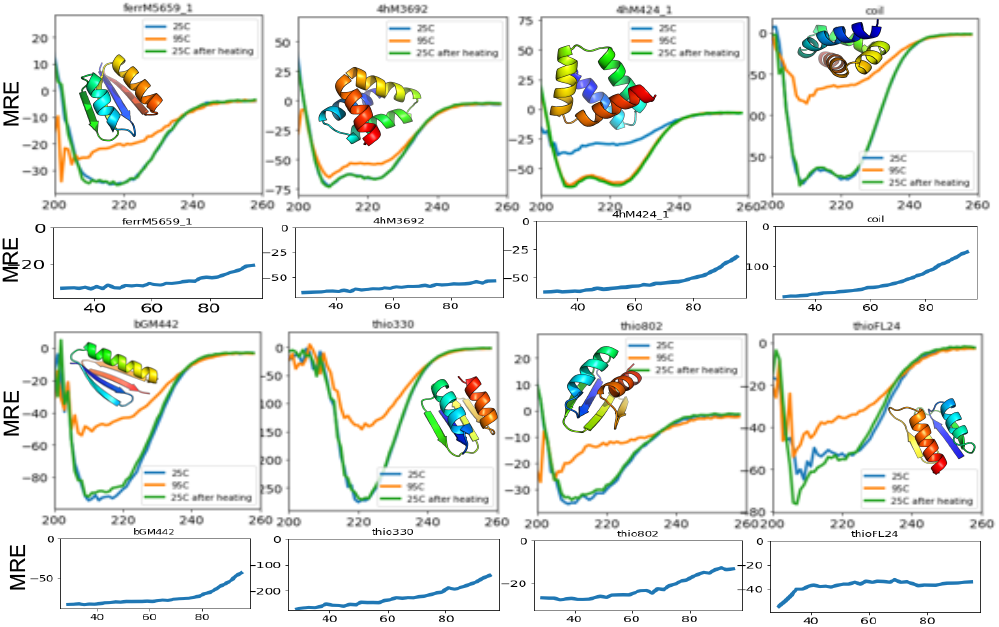
Biochemical and biophysical characterization of a subset of the designed proteins. Circular dichroism spectra measured at 25°C,95°C and 25°C after heating up to 95°C and are in agreement with the expected secondary structure content of the design models. MRE, mean residue ellipticity. Melting curves show MRE at 222 nm at increasing temperatures.

Interestingly, folds varied in their success rates (Fig. 2). Three helical bundles generally showed a higher success rate than any of the other folds; this is consistent with previous report^1^. Surprisingly, the not-seen-in-nature folds F2 and F4 folds had higher success rates than the naturally occurring folds that contained alpha and beta-strands; while thioredoxin and the super helical-coiled four helical bundles (coil) had the lowest. The coil designs alternate between shorter and longer helices while having specific distance constraints to guide supercoiling. Fewer secondary structure contacts and a smaller core when compared to the three helical bundles likely increase the room for error.

So far, the *de novo* design of a thioredoxin fold has not been reported. Here, we attempted the even more challenging problem of minimalizing the overall fold. The thioredoxin fold can be subdivided into an N-terminal *βαβ* motif and a C-terminal *ββα* motif (Fig. 4A), which is commonly connected by a small helix (*α0* or *α2*, Fig.4A,B); the *βαβ* element (characteristic for the alpha/beta family) is found in many larger proteins as it is the connecting motif that enables the expansion of the protein domain space^2^. Its incorporation will generally allow for extensions of the fold and thereby provides means to build larger proteins. Hence, the ability to design this element with high shape diversity provides a tool for building larger or more shape-diverse folds and domains. Commonly, the thioredoxin fold has a three-layer *β/α* sandwich with the central sheet formed by five strands flanked by two *α*-helices on each side. However, many of thioredoxin-like proteins have variations in their *α*-helices or the fifth *β-*strand (Fig. 4A,B). We designed a minimal version of the thioredoxin fold, containing only the core 4 sheets and 2 parallel or anti-parallel helices, replacing the common *α2* helix with an extended loop (Fig. 4G). We solved the structure of one of our designs using nuclear magnetic resonance (NMR, Fig. 4D, S9, S10, Table S2). The NMR ensemble agrees with the designed model with a 1.64 Å rmsd difference that derives mostly from the last helix. All *β-*strands were very close as they were designed (0.9 Å C-alpha rmsd) which therefore present the first accurately *de novo* designed thioredoxin fold.

**Figure 4.**
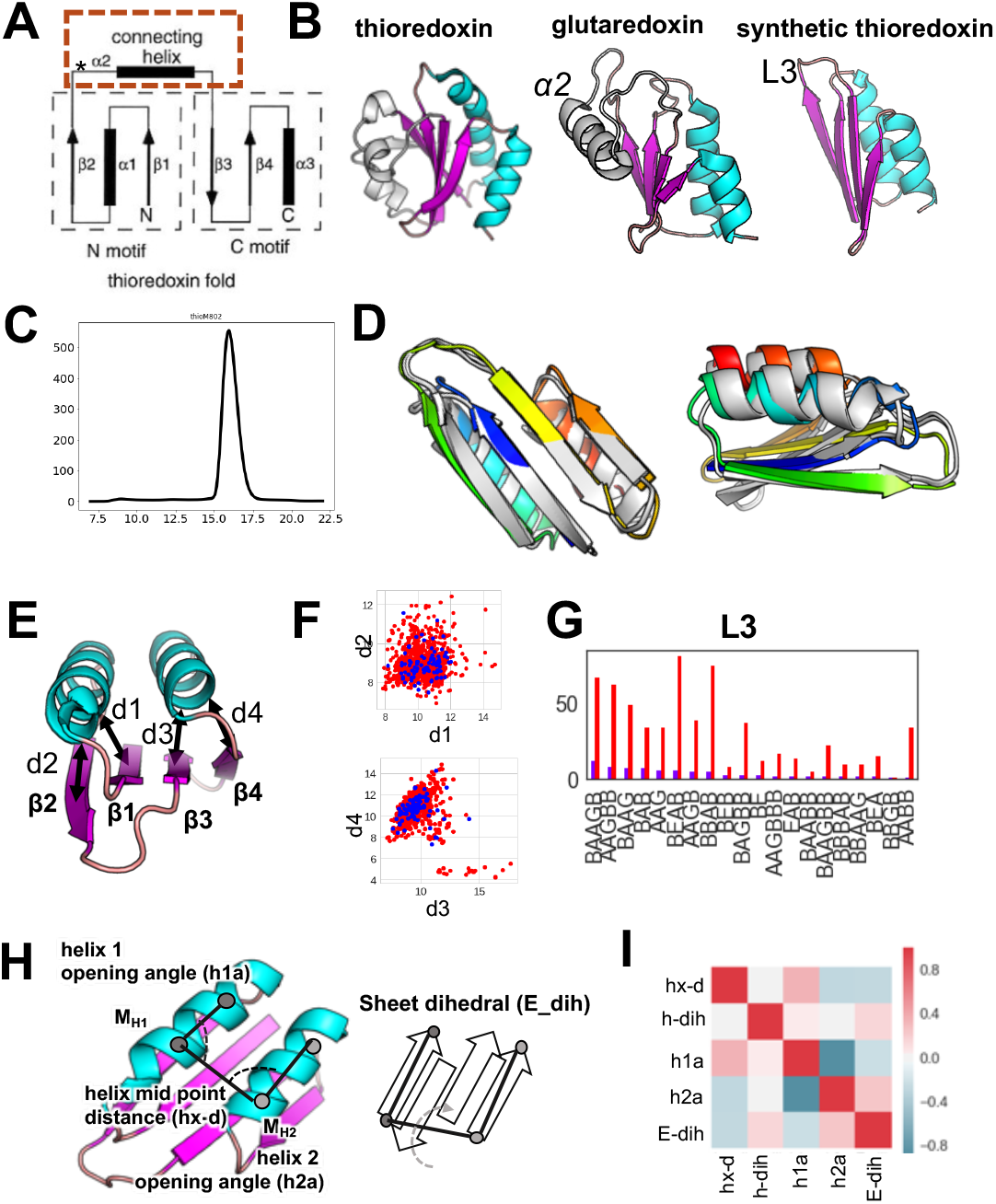
(**A**) Connectivity of the basic thioredoxin fold showing N and C motif; where the N motif represent the domain extending of fold expanding *βαβ* motif. The commonly found *α*0 or *α*2 was replaced for a minimized thioredoxin fold. (**B**) Thioredoxin folds found in nature and the here designed synthetic versions. (**C**) SEC of ems_thioM_802 shows single defined peak using a superdex S75). (**D**) Lowest energy NMR structure (grey) compared to the the model of ems_thio-802. (**E)** Distances measured between secondary structure elements and (**F**) their variations found in stable (blue) and unstable (red) designs. (**G**) Loop variations of the connecting loop L3 which replaces the *α*2 helix of natural thioredoxins, several conformations allow a stable protein. (**H**) Definition of geometric descriptions measured to evaluate shape diversity, including how the sheet dihedral were calculated (**I**) interdependence of structural factors reveal several correlations within a fold.

To explore small alpha+beta proteins, we sampled variations of the beta-grasp (Fig. S11) and extensively variations of the ferredoxin (Fig. S12) fold. Previous attempts to design very small versions of the ferredoxin fold have failed, resulting in unfolded proteins^4^. We extensively sampled secondary structure lengths and registers for ferredoxin fold and found that there are indeed geometric limitations (Fig. S12), likely due to the inherent requirement of right-handed strand-helix-strand motifs. However, we were able to design small and stable variations of this fold of as few as 55 amino acids in length (Fig. S12E).

Parametrically *de novo* designed helical bundles^13^ or repeat proteins^14^ are highly stable. However, the non-parametric small helical bundle fold space can provide much more diverse shapes for new molecular functions. These topologies, particularly four helical bundles (4H), have not been extensively sampled. Furthermore, protein design rules for the loop connections and angle variations are missing; having revealed these with our three and four helical bundles (3H), we implemented these connector rules to our design platform (Fig. S13, S14). To further test our design algorithm, we explored beyond naturally occurring alpha-beta folds. We derived the topologies that have *αββββα* and *βααβββ*, named F2 and F4 fold (Fig. S1). F2/4 designs expressed well and showed a distinct peak using SEC. However, the designs appeared to be dimeric (Fig.S8). Likely, the presence of interconnectedness in a fold, such the *β*3 strand swap to connect the two motifs of the thioredoxin fold, biases against an opening of the structure to form dimers. Folds that have a directly linear connection of secondary structure elements may require further optimization through disulfide connections or perhaps negative design elements that discriminate against a swapped dimeric conformation. However, our pipeline was readily able to produce models for these folds that appeared stable against proteolysis.

To evaluate shape diversity, we compared both protease-stable and unstable proteins for each fold in distances between secondary structure elements, register shifts, dihedral of the outer beta strands to describe curvature of the sheets, dihedrals between adjacent secondary structure elements, interhelix distance, and helix angles given specific *phi* and *psi* angles in their loops, demonstrating that each fold has local plasticity and validating our basic rules of *de novo* protein design (Fig.2, S11-15).

Having a large and diverse set of stable and unstable proteins of different folds available with varying shapes and physical properties, we were able to re-evaluate stability-defining features and develop a classifier based on a Random Forest model that determined the stability of a given fold with high accuracy. In addition, to previously described physical and statistical features for stability, we evaluated additional features describing residue interaction networks and energetic contributions of individual amino acids within tightly connected hubs of residues, resulting in a total of 110 sequence- and structure-based features. The most predictive of the newly introduced features, was the overall energy contributions of the most connected residue: meaning residues that contact many other residues are interaction hubs, they tend to be generally highly buried and likely provide the “glue” of the hydrophobic core. Hence, favorable energetics of these “hub” residues is essential for the protein core formation. Potential clashes likely will result in instability. However, in terms of predictive power we observed as previously seen, that correct local geometry as measured through alignment with short structural fragments (Fig. S16) is the most predictive feature for the folding of a *de novo* designed protein, followed by the number of hydrophobic residues within the protein core. However, unlike previously, by using our larger and more diverse fold data set and descriptive features, we were able to train a model on multiple folds, instead of one-fold at a time^1^ and could predict stabilities of even unseen folds (Fig. 5C). We believe that our more diverse scaffold set is enabling us to learn more general descriptions of these folds and increases the predictive power of the model.

**Figure 5.**
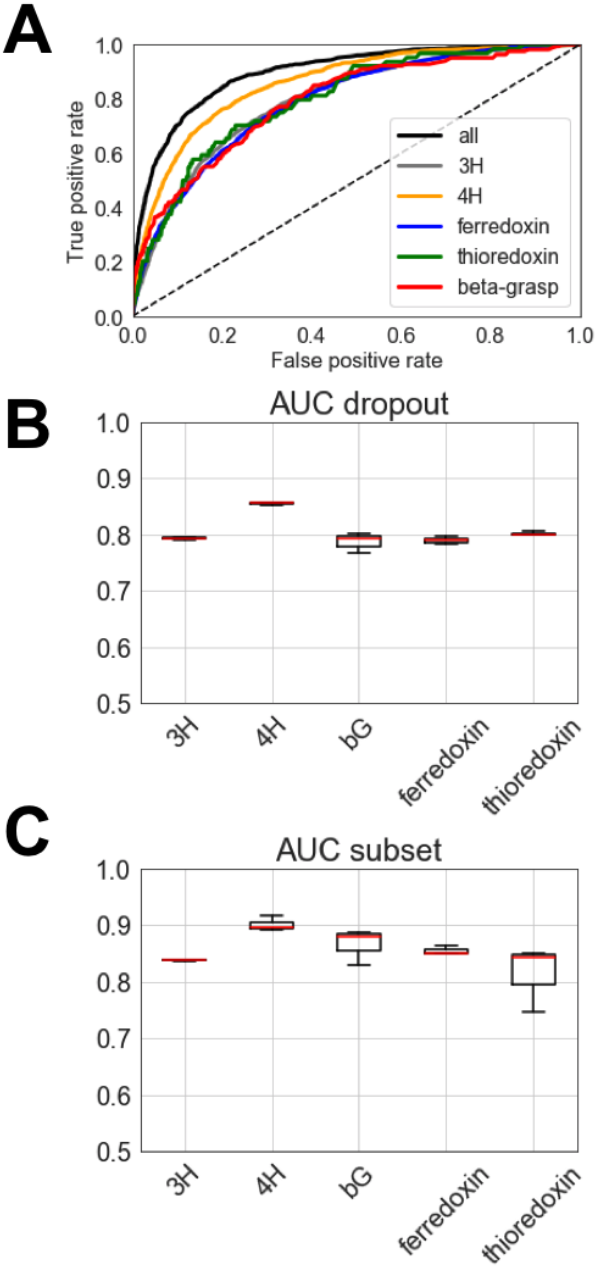
(**A**) Receiver operator curve (ROC) illustrating the predictive power of our Random Forest classifier. To avoid potential bias, we compared the ROC of predicting stabilities either for any protein within the whole set vs a ROC of the different folds after training on the other folds within the set (**B**) AUCs of prediction of an unseen fold; for this prediction, one fold at a time was omitted, and the classifier was trained on all other folds to predict stability of the unseen fold. (**C**) Summary of AUCs of predictions for individual folds.

## Discussion

While previous work simulated the space of a handful of folds^4^ using the elaborate blueprint-based protocols combined with manually defined multi-step assemblies of larger folds, only a few designs were experimentally verified in solution, with the exception of the very short mini-proteins that were examined in context of the development of the protease-based high throughput screen. We built upon previously discovered protein design rules^3, 4^, including connection rules for strands and helices, and provide a versatile fold assembly and design pipeline that allows dynamic sampling of a given fold during the *in silico* folding trajectory. Our extensive sampling and high throughput evaluation allowed us to examine thousands of designs at once revealing geometric diversity of different folds while also exposing more rules for protein design, such as rule to connect helical elements; which we in turn incorporated into the design algorithm. Lastly, our extensive study allowed us to develop a simple prediction model that will help future design approaches to identify stable proteins. The algorithm is implemented into the RosettaScripts^15^ framework, which enables all design features and protocols to be accessed and executed in form of XML files without prior programming knowledge. This work provides a new scaffold database as a general resource for alternative scaffold engineering and protein design projects which recently resulted in the development of picomolar Covid-19 inhibitors^16^.

## Methods

### Requirements for Software and code availability

The Rosetta license can be obtained through Rosetta Commons. For executing the python dependent scripts, PyRosetta and the Anaconda2 package are required. Scripts, xml files and examples will be placed under https://github.com/strauchlab/scaffold_design.

### Computational protein design

#### Overview

Proteins were designed using three steps. First, the backbones were constructed which outlay the three-dimensional structure of the final fold. This step made extensive use of the here developed pipeline and differs from previous approach that have utilized the blueprint builder. The second step involves sequence design for which we utilized two different protocols followed by a last step for the selection of designs to test.

#### Backbone design

The underlying algorithm of the Fold Architect (FoldArchitectMover) has several modular components that work together to design a *de novo* peptide backbone for a fold. Together, these components provide the framework to take a fold-level description (e.g. 3 helix bundle, each helix of length 10-15 a.a. connected by loops of length 3-4 a.a.) and produce protein backbone with the desired secondary structure, realistic geometries, and helix pairing interactions. These modules are part of different “sub-architects,” which uses a user-provided (via XML) description of a fold or subset of residues within a fold to create a set of per-residue instructions; a “pose folder” that applies an *in silico* folding algorithm, which uses the instructions provided by the architect to perform the folding process; a set of “filters,” which can be any Filter recognized by Rosetta, which evaluate the backbone generated by the pose folder to ensure that it is correctly folded; and a “perturber”, which uses an architect to generate a new set of instructions for another folding attempt in the event that the filters did not accept the backbone. Each of these components can be arbitrarily extended to support new algorithms by creating subclasses of the Architect, PoseFolder, Filter, and Perturber classes.

One complication with the *de novo* extended chain folding of backbones is that is scales poorly with length; a single missing hydrogen bond in a backbone can lead to incorrect secondary structure and failure of filters. While a helix of length 15 might be correctly formed after most fragment insertion monte carlo trajectories, a complete 40-50 a.a. miniprotein fold might require thousands of trajectories from extended chain to find one that is correctly folded. This has been addressed in previous work with single folds through extensive intervention by an expert. To address this problem for arbitrary folds, we developed an algorithm (DivideAndConqueror) that identifies subsegments of a full-length backbone that can be folded individually and generates a strategy to build the backbone incrementally, piece-by-piece. Generally, the DivideAndConqueror algorithm uses the architect(s) to split the work for a full backbone into subsegments as small as possible that contain a complete pairing; all possible divisions are considered. For example, for a simple fold with topology EEHE (three antiparallel strands + one helix that is paired to the strands), the algorithm might divide the work by first folding EE (contains a pairing between the two strands), then adding H (contains a pairing between the helix and already-folded strands), and finally E (contains pairing to an already-folded strand). Each subsegment is folded with the modules described above, and once built, is evaluated using the filters. If the model passes the filters, the algorithm then folds the next structural element; if the model fails the filters, the perturber instructs the architects to permute the parameters of the subsegment (e.g. secondary structure lengths, register shift, *phi* and *psi* angles, loop connectivity) and another folding attempt is made. In this way the FoldArchitect gradually adds and folds a backbone while samples different lengths, ABEG0 combinations and other parameters that fit within a combination of user-defined restraints, as well as a series of previously discovered protein design principles to find parameters required for correct properties of the fold.

As there are 5 distinct structure-related areas of *phi*-*psi* angle distributions, proteins can be described with a five letter alphabet^6^: ABEG0 (Figure 1C). This extends the secondary structure description beyond the simple alpha-helical (A), beta-sheet (B) and loop region, yet narrows the description of the loop conformational space which is necessary to guide the *in silico* folding process. Design protocols up-to this point were limited to using a single linear sequence in form of its ABEG0 sequence as a “blueprint” file or possibly a handful of blueprints which we eliminate with our approach in favor of an abstract description of a fold.

Folds were *in silico* folded by inserting structured fragments curated from the PDB^6, 7^ (Fig. S1) specified by the architect-determined ABEG0 sequence into an extended chain of a poly-valine residues.

#### Loop connection sampling

Instead of sampling all possible loop conformation for a given loop length, only loop conformations commonly found between αβ, βα and ββ connections as previously discovered^4^ were sampled. In addition, our study identified new rules for the connection of helical elements which we also incorporated. As more principles are identified, they can be readily added. Furthermore, we enabled the possibilities to provide distance constraints between adjacent elements – if desired, incorporate “bulges” to introduce sheet curvature and *de novo* folding which we took advantage of to build beta-grasps. After defining each of these parameters, the protein is “folded” *in silico* in segments, starting with 2 secondary structure elements with values of the parameters above chosen from the allowed set. If the attempts fail to fulfill all specified filters, new parameters are selected. If the *in silico* folding attempt succeeds, parameters are stored and new cycle to add the next segment will be attempted. Previously discovered protein design priniciples^8^ are respected and the protein is dynamically assembled adding one segment at a time.

#### Pairings

Secondary structure elements that interact with one another in the desired fold are identified by “Pairings.” Movers and filters can then also use this information to obtain information about the desired fold. The different pairing types are “HelixPairing”, which describes a pairing between two helices (e.g. parallel/antiparallel); and “StrandPairing”, which describes a strand-strand pairing (e.g. parallel/anti-parallel, register shift); and “HelixSheetPairing”, which describes an interaction between a helix and a beta-sheet.

#### BetaSheetArchitect

The BetaSheetArchitect is used to define *de novo* beta sheets by combining information from StrandArchitects. Sheets are defined spatially by looking at the face of the sheet (this does NOT use N-->C ordering). Strands are assumed to be paired to the strands that are defined above/below. The architect automatically adds the appropriate strand pairings.

The architect will only attempt to build valid sheets, those where the fully built sheet has no unpaired residues.

#### Distance constraints

Distance constraints of any kind can be applied ambiguously without knowledge of the final length used in the dynamic algorithm. In addition, the user can define different kinds of constraints. Details are within each XML file for the computed folds.

#### Helix “kink”

This filter monitors the curvature of helices and allows to restrict them. We generally only allowed bending of less than 15°.

#### Backbone design

Briefly, each protein secondary structure can be described as a sequence describing the bins of the phi-psi angles in the Ramachandran plot; which we categorize into 5 different bins termed ABEG0 (Fig.1). To build the tertiary structure of a protein, short structured fragments with the desired ABEG0 sequence are used for its assembly. Hence, previously to describe the tertiary structure to be built, the ABEG0 sequence is recorded in an individual blueprint file, which allowed no variation in lengths of any secondary structure element, registers shifts, bulges, as well as distant constraints. The latter would have to be specified for residues in specific blueprint. Further, structures that are longer than 40 amino acids would require two independent folding steps to avoid large amounts of time spent at sampling the possible backbone conformations within a given ABEG0 space. Thereby the infrastructure was limiting to large scale design of highly diverse scaffolds of a desired fold.

For the design of a single protein of a given fold, parameters were defined, including which subsegments are folded initially. Folding of the backbone was followed as previously reported through fragment insertion, with the difference that fold a subsegment at a time, which enables efficient sampling within the pre-defined parameter space. The Perturber adjust dynamically the folding process (for instance by varying lengths or type of loop insertion, register shift) to satisfy all specified parameters of the folded backbone.

#### Sequence design

Backbone constructs representing the complete fold were stringently filtered for the omega and *rama* angles before designing their sequence. Two different design protocols were utilized, one utilizing the previously described pair-motifs^1^ to design the core of the proteins. The pair motif database contains two directly interacting side chains of two amino acids extracted from crystal structures, thus describing a “pair”. We observed that using this protocol, efficient sequence design was observed that passed all filters. The later were local structural geometry, average degree of connectivity within a certain radius to ensure good packing, Rosetta scores and several other(XML code and summary).

### Scoring matrix and design selection

All score terms for filtering and evaluating designs are summarized in the SI. Their implementation can be found in the design protocols sequencedesign.xml and sequencedesign_w_motifs.xml, rescore15.xml and rescore16.xml followed by adding terms previously reported as “enhanced_score.sc”^1^. Resulting in 2,000-12,000 finished design models per fold.

### Random Forest prediction model

Models were fit using the scikit.learn package. We used 500 as numbers of trees for the estimator and “gini” as criterion while allowing “out-of-bag” samples to estimate the generalization accuracy. While building the trees, default bootstrapping sampling was allowed.

### Library generation

Amino acid sequences of designed proteins were encoded into DNA using DNAworks2.0 and “ecoli2” codons^17^. Oligo libraries encoding designs and control sequences were purchased from Agilent Technologies as part of a 27,000 oligonucelotide pool. Genes shorter than 230, additional amplification sequences were added as previously reported^2^ in order to amplify sequences equally. Amplification was performed using a qPCR (BioRad) to avoid overamplification. The number of cycles was chosen based on a test qPCR run in order to terminate the reaction at 50% maximum yield. Second, this reaction product was gel extracted to isolate the expected length product, and re-amplified by qPCR to obtain larger amounts. The amplified PCR product was gel extracted and concentrated for transformation of EBY100 yeast^18^ (1-2 µg of insert and 1 µg of linearized vector). Yeast display employed the pETCON3^19^ which was linearized by digesting its DNA with *NdeI* and *XhoI*. The amplified libraries included 40bp segments on either end to enable homologous recombination with the pETCON vector. Gel extraction and PCR purification were performed using QIAquick kits (Qiagen Inc).

### Yeast display proteolysis

Protease reagents Trypsin-EDTA (0.25%) solution was purchased from Life Technologies and stored at stock concentration (2.5 mg/mL) at −20°C. α-Chymotrypsin from bovine pancreas was purchased from Sigma-Aldrich as lyophilized powder and stored at 1 mg/mL in TBS +100 mM CaCl2 at −20°C. Each reaction used a freshly thawed aliquot of protease.

EBY100 yeast cell cultures were induced for 16-18 h at 30°C in SGCAA^20^. Induced cells were digested with increasing concentrations of chymotrypsin and trypsin in separate tubes. Cells were normalized to 1 mL at O.D. 1 (12-15M cells), washed and resuspended in 250 μL buffer (20 mM NaPi 150 mM NaCl pH 7.4 (PBS) for trypsin reactions, or 20 mM Tris 100 mM NaCl pH 8.0 (TBS) for chymotrypsin reactions). Protolysis was initiated by adding 250 μL of room temperature protease in buffer (PBS or TBS) followed by vortexing and incubating the reaction at room temperature (proteolysis reactions took place at cell O.D. 2).

The library was assayed at five protease concentrations over different rounds of sequential selection rounds as summarized in the experiments.csv file. For trypsin digestions we used 0.07 μM, 0.21 μM, 0.64 μM, 1.93 μM, and 5.78 μM protease; chymotrypsin assays used 0.08 μM, 0.25 μM, 0.74 μM, 2.22 μM, and 6.67 μM protease. Selections at lower concentrations (selection strength 1 −3) concentrations of each protease were performed starting from the freshly transformed and induced yeast library. The following these selections, higher concentration conditions were performed as indicated in the *experiment*.*csv* file. This file contains the precise order of selections, including cells sorted and selected. Selection strengths (as indicated under “parent” reflects above listed concentrations respectively; parent 0 represents the starting library pool, whereas 1 reflects the lowest concentration as listed above (0.07 μM for trypsin and 0.08 μM for chymotrypsin. The highest concentration of protease was parent 5.78 μM for trypsin indicated as selection strength 5. These data are included in the file *experiments*.*csv*, and were used in the EC_50_ fitting procedure.

After 5 minutes, the reaction was quenched by adding 1 mL of chilled buffer containing 1% BSA (referred to as PBSF or TBSF), and cells were immediately washed 4x in chilled PBSF or TBSF. Cells were then labeled with anti-c-Myc-FITC for 10 minutes, washed twice with chilled PBSF, and then sorted using a Sony SH800 flow cytometer using “Ultra Purity” settings. Events were initially gated by forward scattering area and back scattering area to collect the main yeast population, and then by forward scattering width and forward scattering height to separate individual and dividing cells (which were used for analysis) from aggregating cells. Following these gates, cells were gated by fluorescence intensity in one-dimension (Fig. 1B). Small adjustments were made to this gate based on daily conditions to maximize the separation between the major displaying and non-displaying populations. For each sort, we recorded the fraction of cells passing the fluorescence threshold before proteolysis (using cells from the same starting yeast population, but untreated with protease) and after proteolysis, and also recorded the total number of cells collected for each condition. Generally, about 10 million cells were sorted for each protease concentration.

### Next generation sequencing and processing of raw deep sequencing data

Plasmids of sorted and unsorted populations where extracted using the Zymo prep kits (Zymo, version 2) with modifications as described previously^19^. Briefly after DNA extraction the prep was digested with Exo1 and Lambda exonuclease (NEB). Cells were frozen at −80°C before and after the zymolase digestion step to promote efficient lysis. One-half the plasmid yield from the Zymoprep was used as the template for the first PCR amplification. Illumina adapters and 6-bp pool-specific barcodes were added in the second qPCR step. Unlike libraries prepared for transformation, DNA prepared for deep sequencing was gel extracted following the second amplification step. The DNA was pooled and sequenced using a mid-size kit on a NextSeq (Illumina) sequencer. Each library in a sequencing run was identified via a unique 6 bp barcode. Following sequencing, reads were paired using the PEAR program^21^. Reads were considered counts for a particular ordered sequence if the read (1) contained the complete *NdeI* cut site sequence immediately upstream from the ordered sequence, (2) contained the complete *XhoI* cut site sequence immediately downstream from the ordered sequence, and (3) matched the ordered sequence at the amino acid level.

### EC_50_ estimation from sequencing counts

To determine protease resistance from our raw sequencing data we used the previously reported probabilistic model to calculate maximum a *posteriori* estimates of the protease EC_50_ of each member of the pool. It assumes that proteolysis (i.e. any cleavage that results in detachment of the epitope tag) follows pseudo-first order kinetics, with a rate constant specific to each sequence. Scripts were used exactly as previously reported without modification^1^ and directly taken from the reported repository https://github.com/asford/protease_experimental_analysis.

### Expression of individual proteins, purification and characterization

Genes for selected design for detailed biochemical evaluations were cloned into pET29b+ and expressed in Lemo21 cells (DE3) (NEB) supplemented with 50 ug/mL kanamycin either using Studier autoinduction media^22^ or at 18°C in Terrific Broth (TB) media using 1 mM IPTG for a 3-4 h induction at an O.D. of 0.7. Briefly, starter cultures were grown overnight at 37°C TB medium overnight with added antibiotic and used to start a 500 mL culture at a 1/50 dilution. His-tagged proteins were purified using a nickel column purification step (QIAGEN). Following IMAC, designs (labeled and unlabeled) were further purified by size-exclusion chromatography on ÄKTAxpress (GE Healthcare) using a Superdex 75 10/300 GL column (GE Healthcare) in PBS buffer. The monomeric fraction of each run (typically eluting at the 15 mL mark) was collected and immediately analyzed by CD or flash frozen in liquid N2 for later analysis. Circular dichroism Far-ultraviolet CD measurements were carried out with an AVIV spectrometer, model 420 or an Olis DSM 1000 CD Spectrometer. Wavelength scans were measured from 260 to 195 nm at 25 and 95°C. Temperature melts monitored dichroism signal at 220 nm in steps of 2°C/minute with 30s of equilibration time. Wavelength scans and temperature melts were performed using 0.35 mg/ml protein in PBS buffer (20mM NaPO4, 150mM NaCl, pH 7.4) with a 1 mm path-length cuvette.

Protein concentrations were determined by absorbance at 280 nm measured using a NanoDrop spectrophotometer (Thermo Scientific) using predicted extinction coefficients. Protein concentrations for designs lacking aromatic amino acids were measured by Qubit protein assay (ThermoFisher Scientific).

### Isotope labeling for NMR

The expression of uniformly ^13^C, ^15^N-labeled protein for NMR analysis utilized M9 media with ^13^C glucose and ^15^N ammonium salts (Sigma) as the sole carbon and nitrogen sources, respectively. A 30 mL overnight culture was grown and used to induce a 500 mL M9 culture. At an O.D. of 0.5, cells were induced with 1 mM IPTG and grown overnight at 18C. Purification was performed as described above.

### NMR spectroscopy and solution structure determination

All NMR spectra were acquired using a Varian INOVA instrument operating at 600 MHz (^1^H). The temperature of the sample was maintained at 25 °C. A single sample of uniformly ^13^C, ^15^N-labeled protein was used for all experiments. The included ~0.6 mM protein, 10 mM sodium phosphate, 10 mM NaCl, trace amounts of glycerol, and 5% D_2_O. The pH of the sample was ~6.8. The sample volume was approximately 300 uL in a susceptibility-matched NMR tube (Shigemi Inc.). Chemical shifts were referenced in the recommended manner using an external, standard sample of Na^+^DSS^-^ in D_2_O^23^. Data were processed and analyzed using Felix NMR.

The main chain, and some side chain, chemical shifts were assigned using established triple resonance approaches employing a standard suite of experiments ((^1^H, ^15^N)-HSQC, (^1^H, ^13^C)-HSQC, HNCA, HN(CO)CA, HNCACB, CBCA(CO)NH, HNCO, HN(CA)CO, HBHA(CBCACO)NH). Remaining side chain resonances were assigned using TOCSY, NOE, and aromatic-specific experiments (HCCH-TOCSY, HCCH-COSY, H(CCO)NH-TOCSY, C(CO)NH-TOCSY, (HB)CB(CGCD)HD, ^1^H-TOCSY relayed constant-time (^1^H, ^13^C) HMQC (aromatics), NOESY-(^1^H, ^15^N)-HSQC, NOESY-(^1^H, ^13^C)-HSQC, NOESY (^1^H, ^13^C)-HSQC (aromatics)).

## Data availability

NMR structure has been deposited in the Protein Data Bank under accession code 7LDF. Scripts, xml files and examples will be placed under *https://github.com/strauchlab/scaffold_design*.

## Acknowledgements

EMS was supported by the R01 AI140245, R21 AI143399, and by the Washington Research Foundation. D.B was supported by the Howard Hughes Medical Institute (D.B.). We thank Inna Goreshnik for amplifying the initial library pool and Jeremy Mills for commenting on the manuscript.

## Author contributions

TL wrote the Rosetta-based FoldArchitect code, EMS developed the fold-specific, design protocols and stability prediction model, EMS designed the proteins, KN, AT, RC, LC, JU and EMS performed the experiments, JU solved the NMR structure, EMS and DB conceptualized the work.

