## Supplementary material for "Sampling of Structure and Sequence Space of Small Protein Folds": SI

**Additional files:**

files will also be available under: *https://github.com/strauchlab/scaffold_design*.

### Supplementary figures

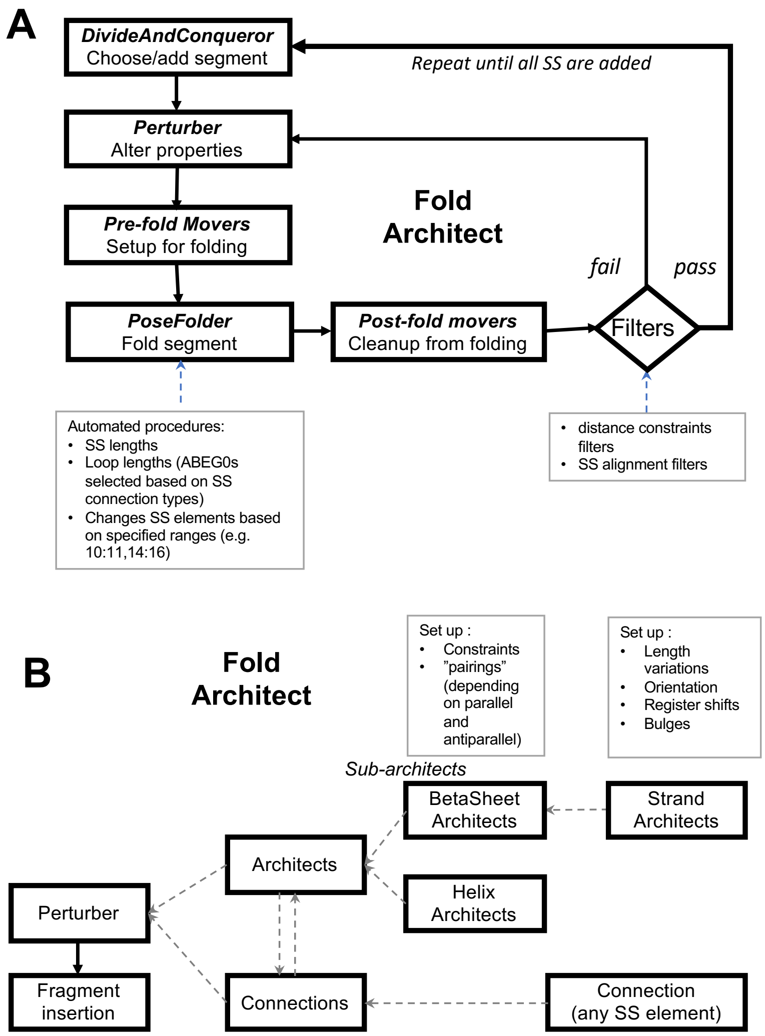

**Figure S1.** Workflow for backbone design.

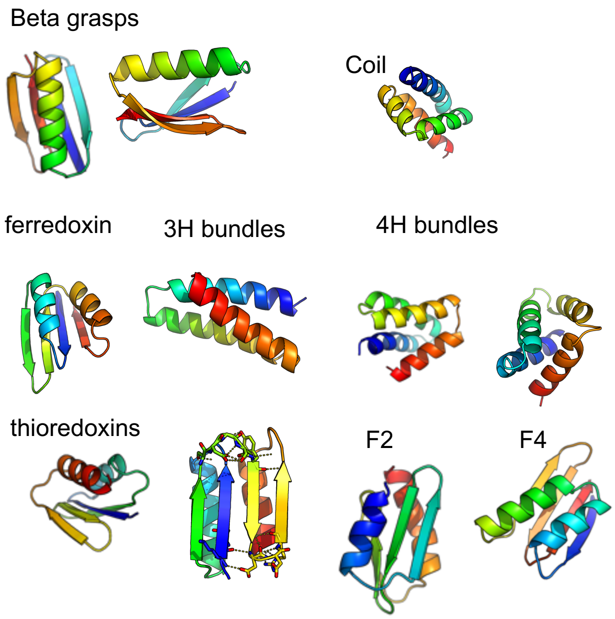

**Figure S2**: Overview of sampled topologies that can be encoded by 64 residues or less.

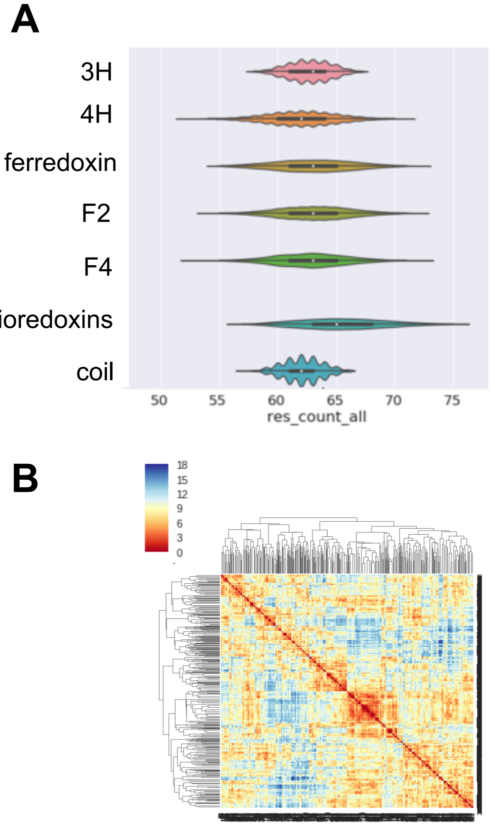

**Figure S3. Sampled diversity for backbone generation.** (**A**) Length histogram of generated backbones using a dynamic assembly of the FoldArchitect. Only backbones that had less than 65 amino acids were subjected to sequence design and subsequentially tested using the protease-based yeast surface display screen. (**B**) Comparison of 3,500 backbone designs for ferredoxins show that the ABEG0 sequences of generated designs are highly diverse. Comparison was computed using Levensthein distances.

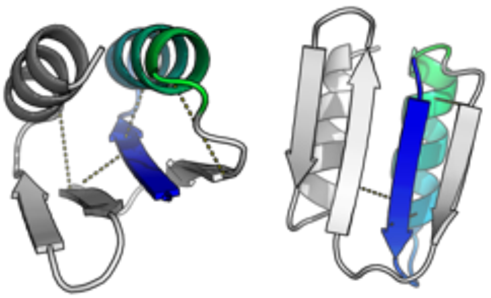

**Figure S4. Starting point and distance constraints used for thioredoxin folds.** For each fold, the two starting segments need to be defined; however the default is to start with the middle segments if the user does not specify. This worked for most folds except for thioredoxin. For thioredoxins, multiple starting points for the assembly were tried. Starting with the first strand and helix resulted in the most decoys passing all filters and was thus used for the backbone generation protocol. Loose distance constraints were defined for each element (dotted yellow lines).

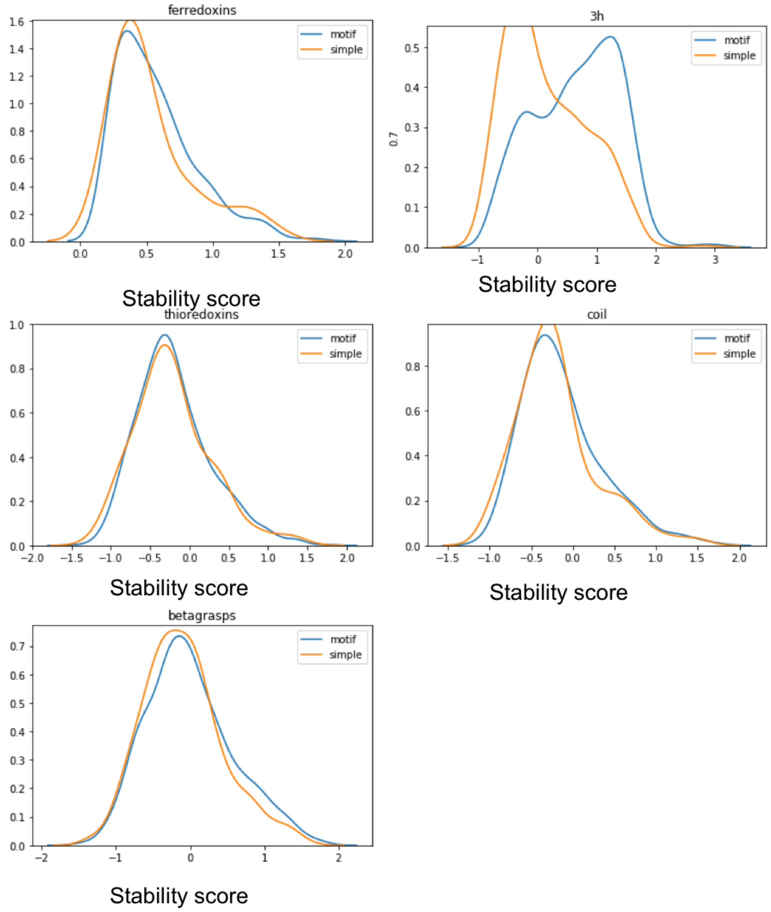

**Figure S5. Difference in stability of different folds depending on their sequence design protocols.** We compared the stability scores for each design and separated two populations based on the design protocol used for sequence design. One was taking advantage of "pair-motifs" before using the Rosetta-based design protocol that is implemented into the Mover FastDesign.

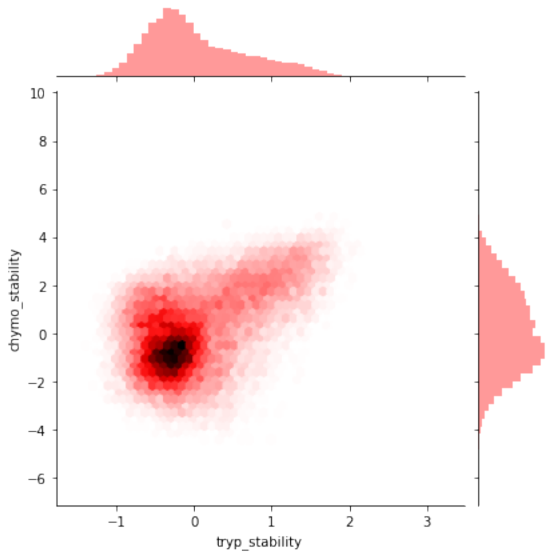

**Figure S6**. Stability scores for chymotrypsin (y-axis) and trypsin (x-axis)

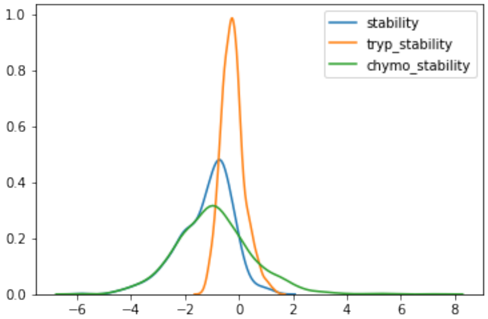

**Figure S7**. Trypsin and chymotrypsin based stability scores and general stability score (by taking the lower stability value) of 2,300 randomly scrambled sequences that were added as a control. Combined information reveals a threshold of 0.5.

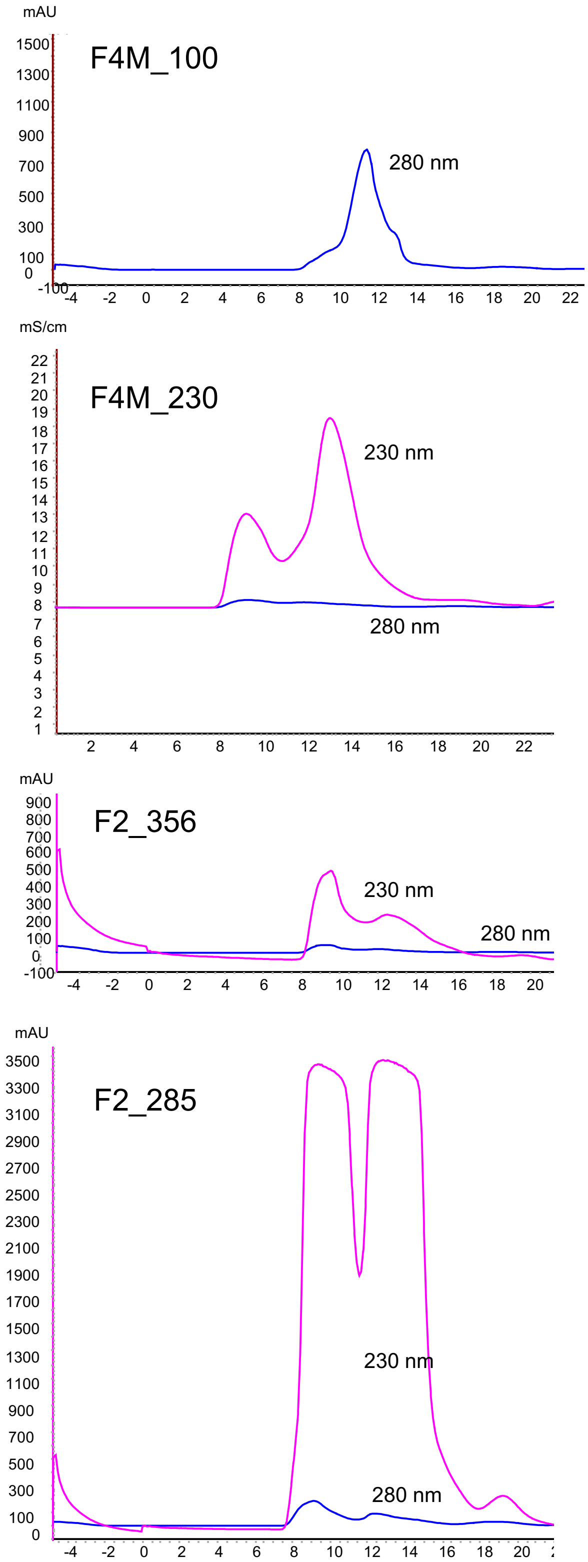

**Figure S8. SEC of F2 and F4.** For proteins without tryptophans, we show the 230 nm wavelength as well.

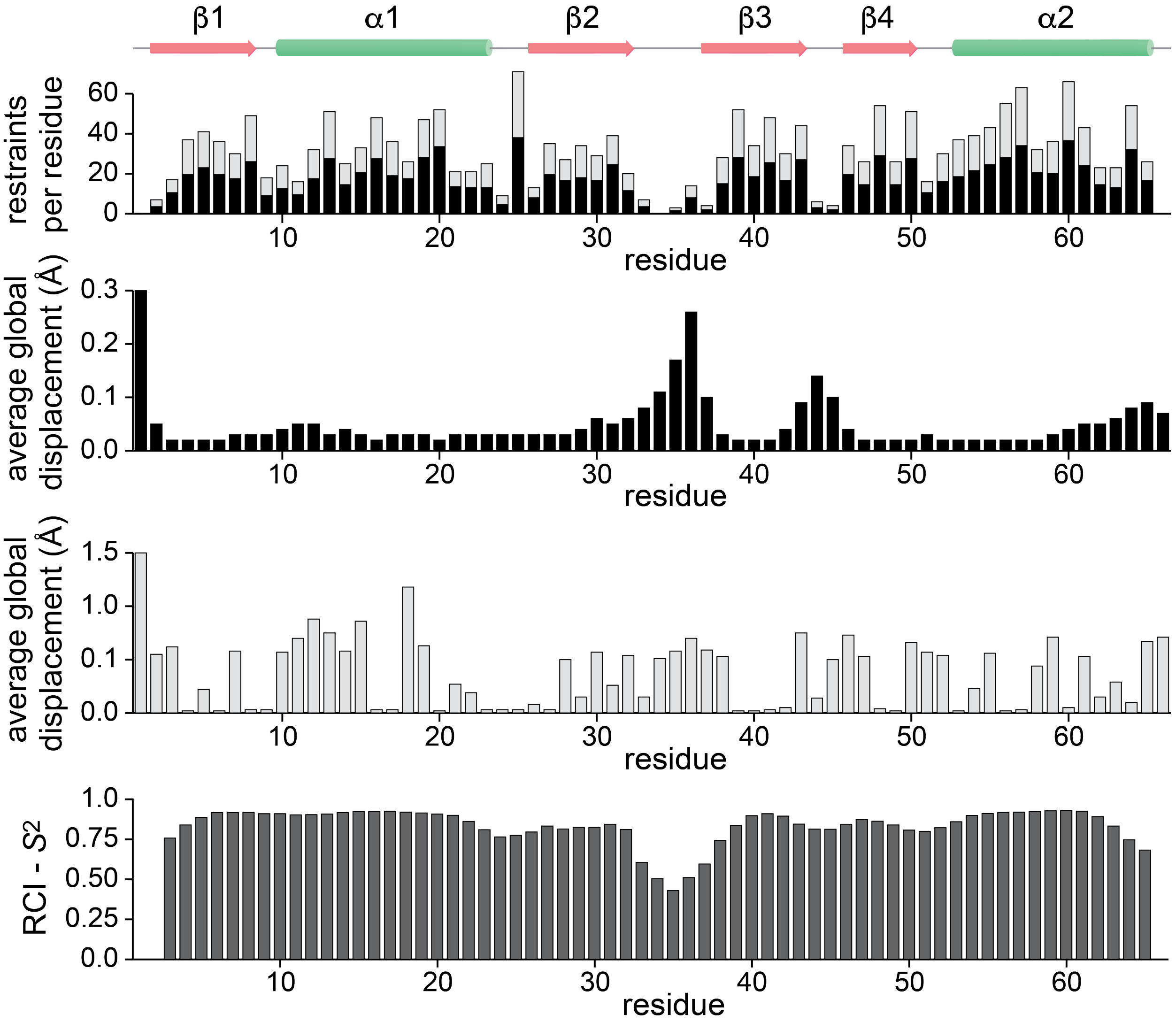

**Figure S9**. Site-resolved distance restraint densities, average distance displacements, and estimated order parameters for the 20 structures in the NMR ensemble. (*Top*) Plotted are the numbers of NOE-derived distance restraints per amino acid residue used in the structure calculations. The height of the black portion of each bar is the number of restraints for that residue calculated as 0.5 for each of the two atoms involved, then summed for each residue. The sum over all residues is then equal to the total number of NOE-based distance restraints (1138, see Table S2). The overall height of each bar (light-colored) is the number of restraints calculated with each restraint counted once for each residue involved (intra-residue restraints count as one for the residue, inter-residue restraints count once for each of the two residues). (*Center*) For each residue in the 20-structure ensemble, the magnitudes of the average global distance displacements (from the mean) for the main chain atoms (black bars, top graph) and heavy atoms (light bars, bottom graph) are shown, in angstroms (Å). The main chain graph is truncated at 0.3 Å (the value for the N-term residue is 0.75 Å), and the heavy atom graph is truncated at 1.5 Å (the value for the N-term residue is 1.57 Å) in order that the values for the other residues can be more easily discerned from the graphs. (*Bottom*) Estimates of the model-free order parameter, $S^2$, based on chemical shifts from random coil index (RCI) calculations.

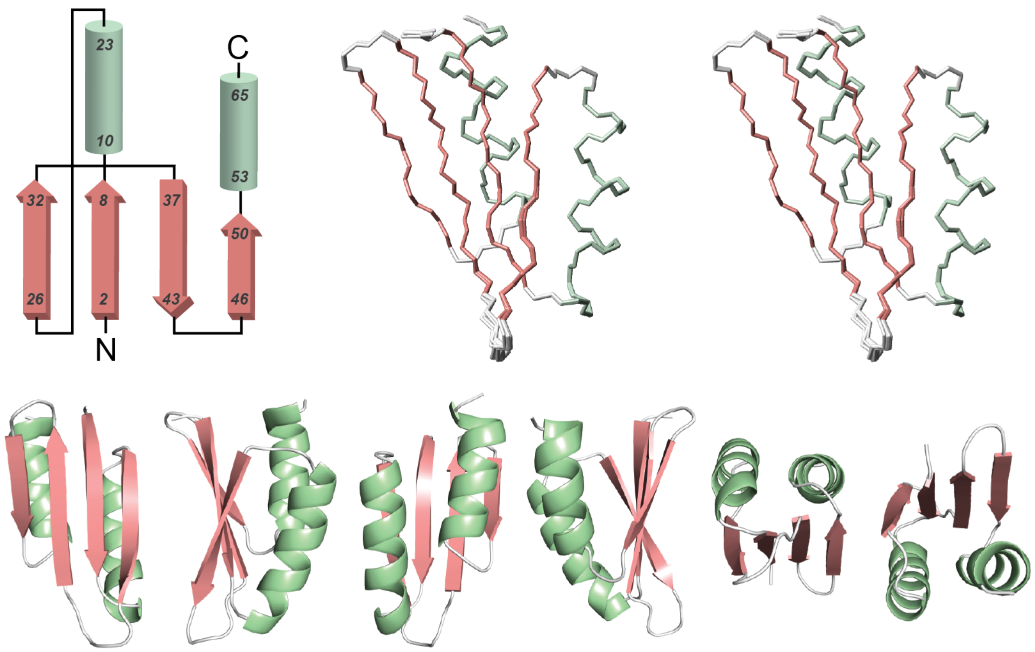

**Figure S10**. Thio-802 structure and topology. (*Top, left*) Topology 'wiring diagram' of the solution structure of the Thio-802 protein determined using NMR spectroscopy (no register information is implied). (*Top, right*) Cross-eyed stereo view of the 20-structure ensemble (refined structures with the lowest overall energies) superimposed on the mean structure. (*Bottom row*) Various views of the ribbon diagram of the lowest energy structure of the ensemble.

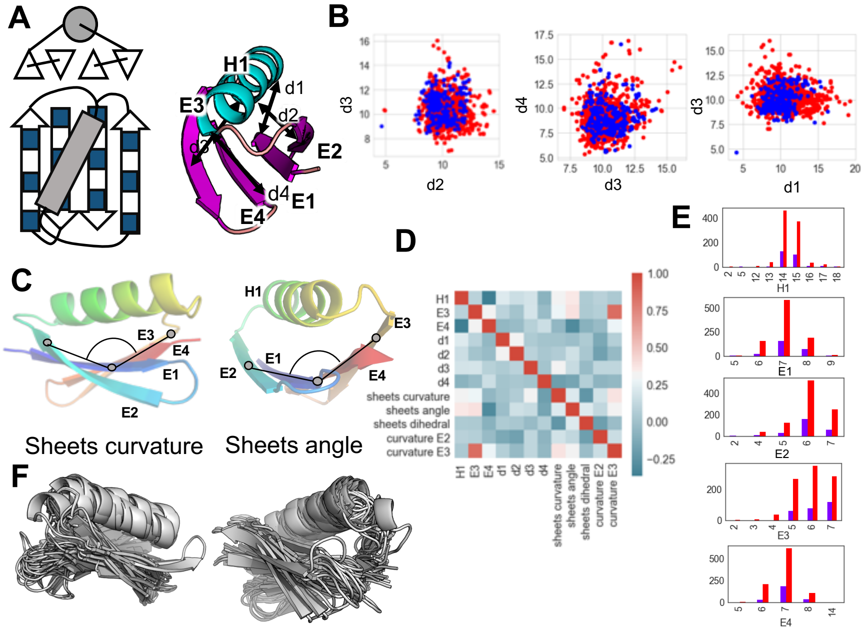

**Figure S11. Geometric analysis of beta-grasps.** (**A**) Schematic of the beta-grasp fold and labeling of distances between midpoints of strands and helix. (**B**) Plots of distances between strands and helix; blue represents stable scaffolds and red instable. (**C**) illustration of how sheet curvature and sheet angle was measured. (**D**) Correlations of distances, specific secondary structure lengths and angles for stable folds. (**E**) Summary of lengths of strands and helix for stable (blue) and instable (red) designs. (**F**) Superposition of 20 designed beta-strand folds to illustrate structural diversity.

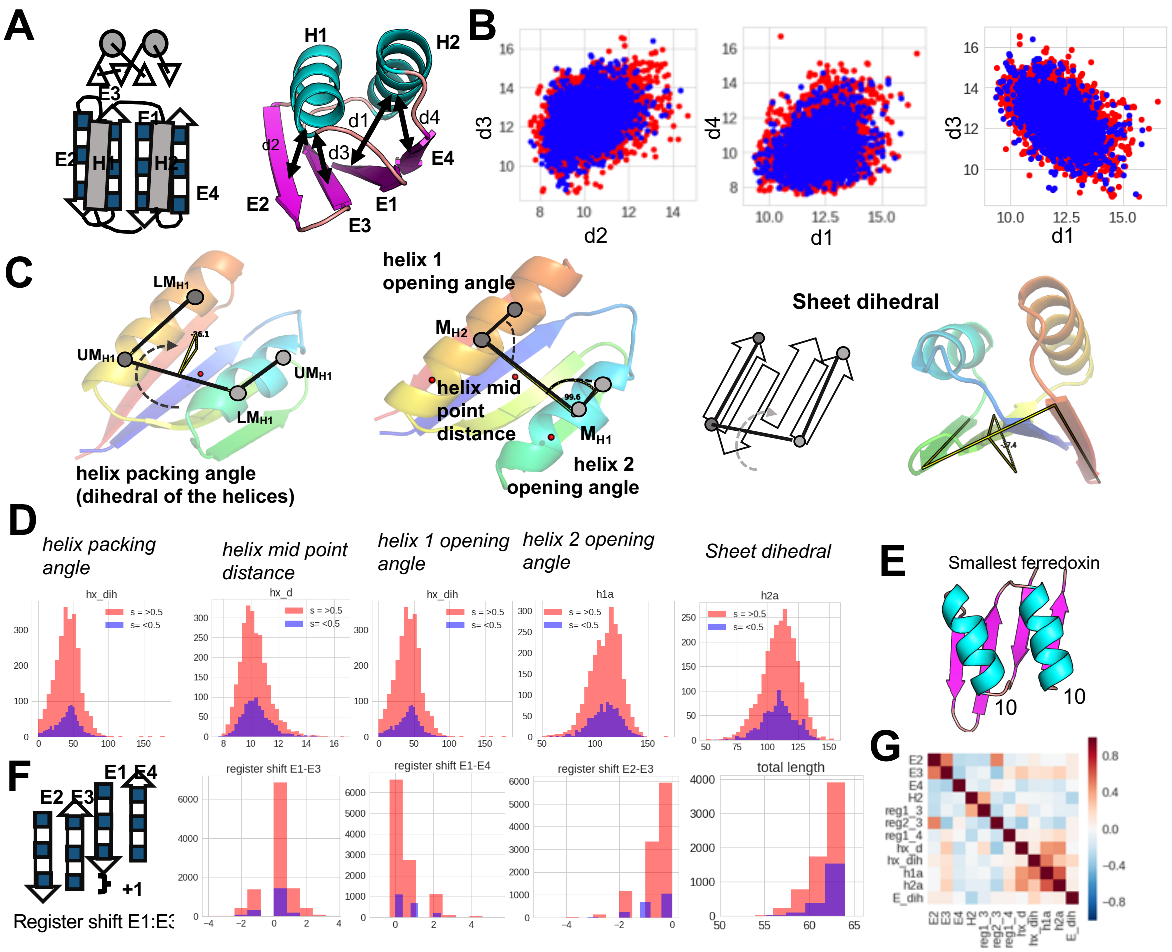

**Figure S12. Geometric analysis of designed and assayed ferredoxins**. (**A**) Schematic of the ferredoxin fold and labeling of distances between midpoints of strands and helices that were measured here. (**B**) Scatter plots of distances between strands and helices; blue represents stable scaffolds and red instable. (**C**) Illustration of how helix distances, angles and sheet dihedral were measured. (**D**) histograms of angles and distances showing stable (blue) and instable (red) designs. (**E)** Model of the smallest, stable ferredoxin sampled; it is 55 residues long. (**F**) Schematic of register shift samples and their distributions. (**G**) Correlations of distances, specific secondary structure lengths, register shifts and angles for stable folds.

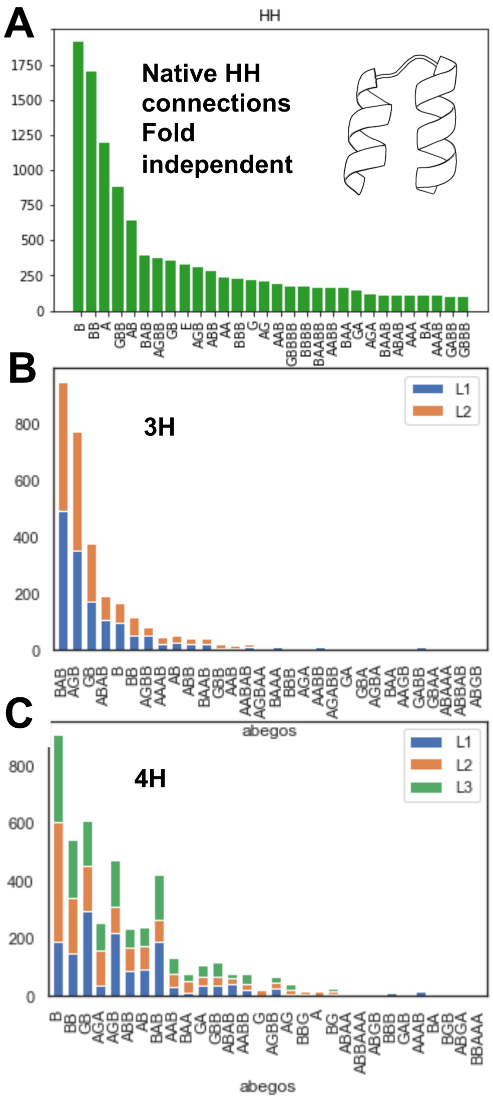

**Figure S13. Rules for connecting helical elements.** (**A**) Distributions of loop connections found in 7000 high resolution crystal structures and (**B**) found in the stable 3-helical (3H) or (**C**) 4-helical bundles (4H) of this work.

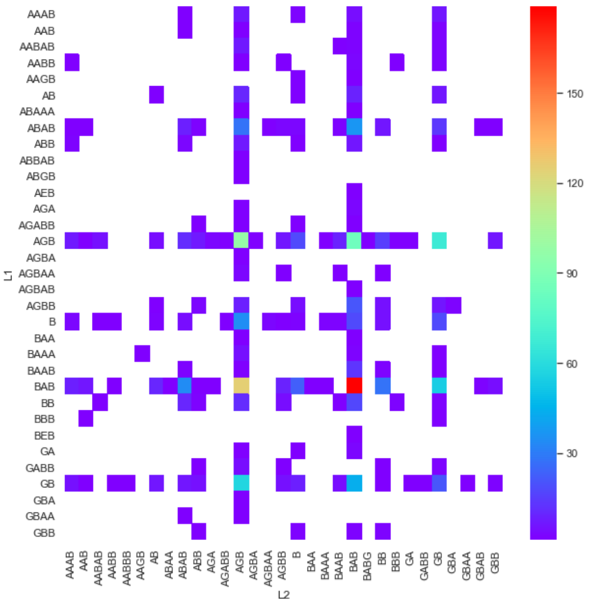

**Figure S14. Loop angles described as ABEG0 letters for stable 3H bundles.** L1 as y-axis and L2 as x-axis. Color describes how often it was seen.

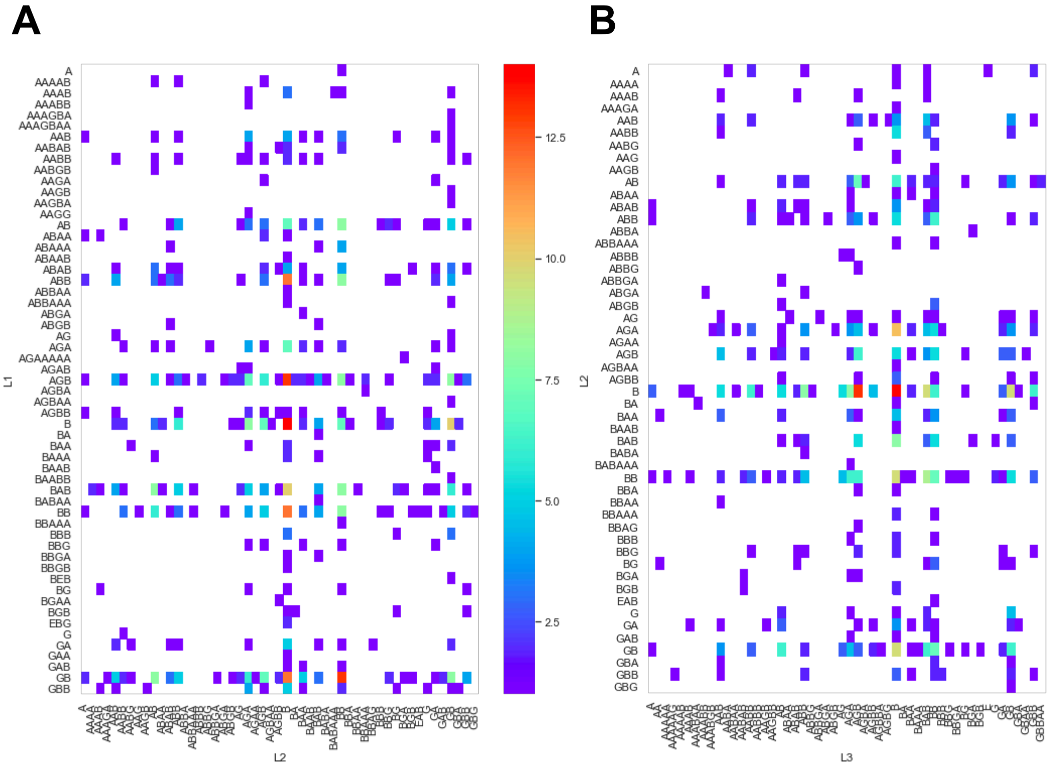

**Figure S15. Loop angles described as ABEG0 letters for stable 4H bundles.** (A) L1 as y-axis and L2 as x-axis. (B) L2 as y-axis and L3 as x-axis. Color describes how often it was seen.

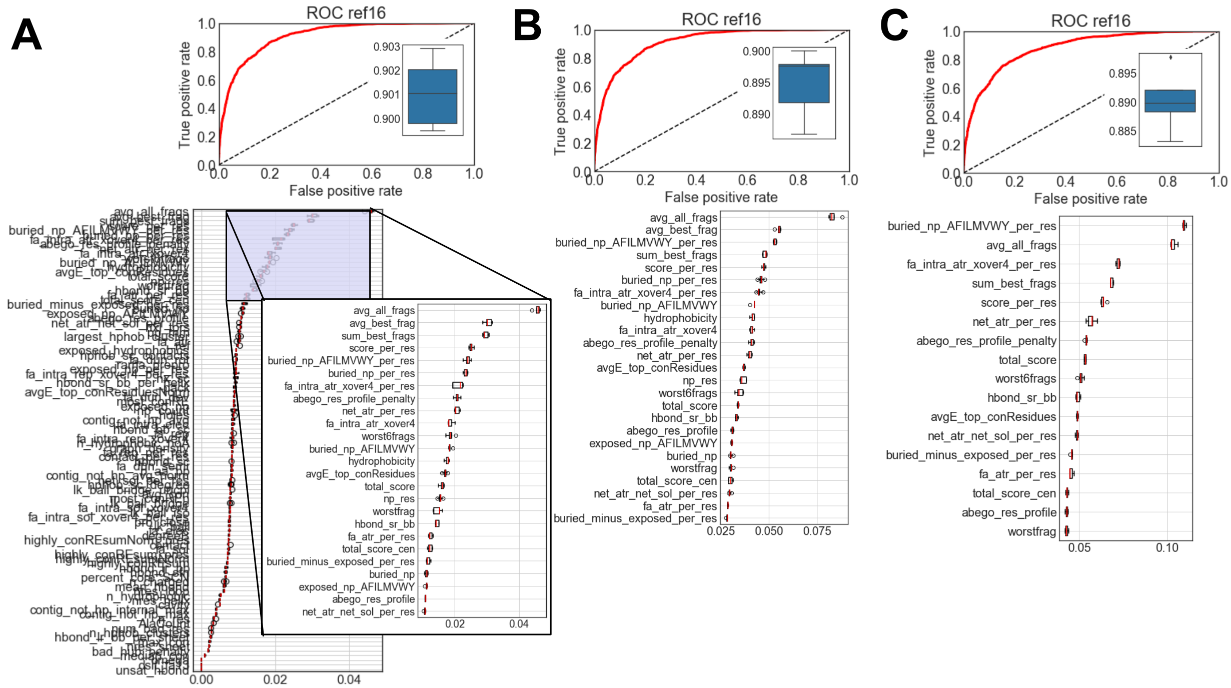

**Figure S16**: **Features of prediction models.** (A) Taking all 110 scoring features, we achieved and AUC of about 0.9 for all designs, using 5-fold cross-validations with 20% of the data. (B) Taking the top 25 determining features to repeat training and prediction; however, the AUC is getting lower and feature importance changes, likely due to correlated terms. (C) After taking out all correlated features, training and prediction was done with 18 features. However, predictions did not improve. Thereby the complete set of features is most descriptive for the stability of these *de novo* designed proteins.

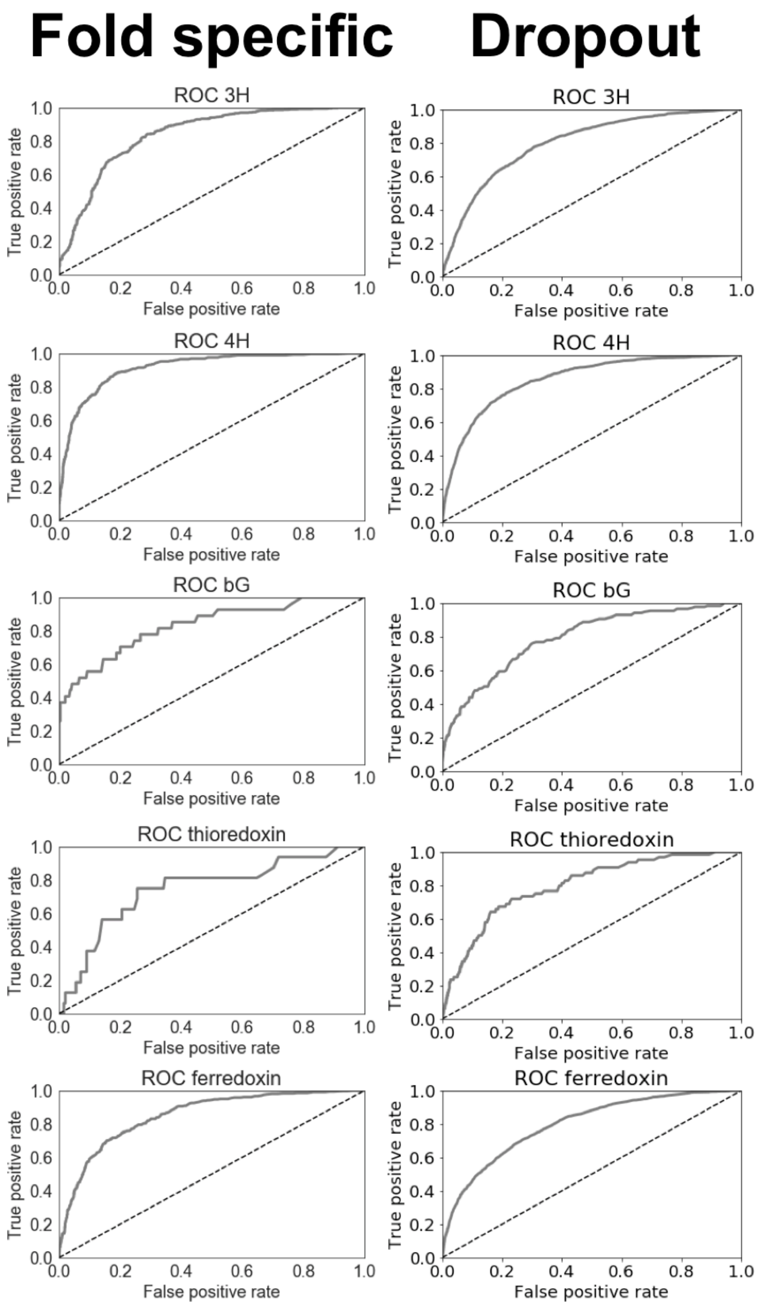

**Figure 17.** ROC for either individual folds or dropouts. The first scenario uses data from a fold, takes 80% of the stability data for the given fold for training and predicts false positives and true positisve of the remaining 20%. For the “dropout” category, all protein sequences and data points are taken out of the complete set. The model is trained on all other folds and FP and TP for the specified fold are then predicted.

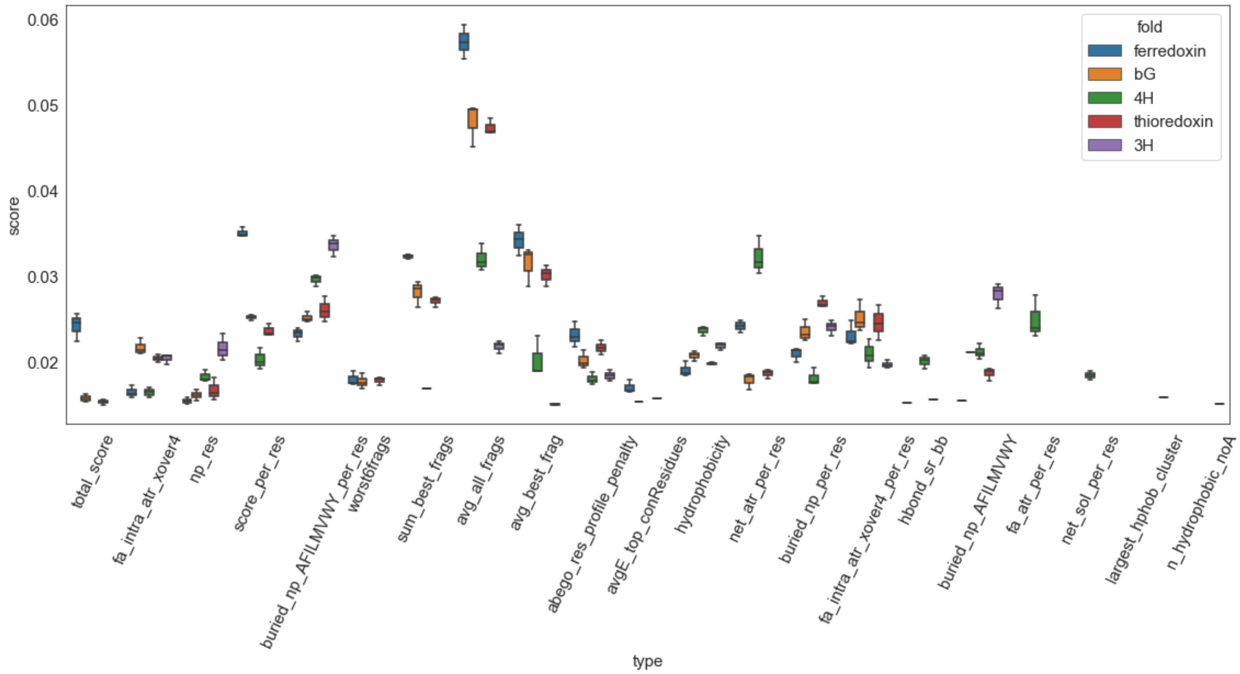

**Figure S18.** Determining features and their contributions to stability predictions dependent on fold. Beta-sheet-containing proteins need an excellent fragment score, where the overall Rosetta score is the most predictive for 3H bundles.

### Supplementary Tables

**Table S1.** Characterization of individual proteins. All proteins with monomeric fractions were folded as measured via CD.

| name | chymo_stability | tryp_stability | stability score | expression yields | oligomeric state (dominant species) |
|---|---|---|---|---|---|
| coil 109.2 | 1.7 | 1.51 | 1.51 | ++++ | **M** |
| bGM_518.1 | 2 | 1.8 | 1.80 | ++++ | A,D, **M** |
| bGM_166 | 1.4 | 1.69 | 1.43 | + | M+other |
| bGM_649.2 | 0.6 | 1.51 | 0.65 | + | **A**, D, M |
| bGM_649 | 0.3 | 0.68 | 0.27 | no expression | N/D |
| bGM_442 | 1.1 | 1.49 | 1.06 | + | M |
| thioFL24 | | | | +++ | **M** |
| thioM_802 | 0.8 | 0.54 | 0.54 | ++++ | M |
| thioFL2_153 | 1.6 | 1.03 | 1.03 | + | **A**, D, M |
| thioM_330 | 1.8 | 1.44 | 1.44 | +++++ | **M** |

| | | | | | |
|---|---|---|---|---|---|
| 4hM_3692 | 1.1 | 3.71 | 1.06 | ++++ | **M** |
| 4hM_424.1 | 1.9 | 3.74 | 1.89 | + | A,D, **M** |
| ferrC_1045 | 1.3 | 1.2 | 1.20 | + | **M** |
| 4hM_4108.2 | 2.3 | 3.82 | 2.27 | + | **M** |
| ferrM_4961 | 1.6 | 1.38 | 1.38 | - | N/D |
| ferrM_5659.1 | 1.4 | 3.8 | 1.45 | ++++ | **M** |
| ems_f2_286 | | | 0.00 | | A, D |
| ems_f2_356 | | | 0.00 | | A, D |
| ems_f4m_100 | | | 0.00 | | **D** |
| ems_f4m_230 | | | 0.00 | | **D** |

**Table S2. Statistics summary for the 20 structure Thio-802 NMR ensemble**

| | |
|---|---|
| **NOE-based distance restraints** | |
| Intra-residue ($i = j$) | 256 |
| Sequential ($\|i-j\| = 1$) | 311 |
| Medium Range ($2 \leq \|i-j\| \geq 4$) | 241 |
| Long Range ($\|i-j\| \geq 5$) | 330 |
| Total | 1138 |
| **Hydrogen bond distance restraints** | 72 |
| **Dihedral angle restraints ($\phi$ and $\psi$)** | 116 |
| **Restraint Violations** | |
| NOE > 0.2 Å | 0 |
| Dihedral angle > 5º | 0 |
| **RMSD from the mean structure** (Å)* | |
| Main chain (residues 2-63) | 0.10 ± 0.04 (0.09 ± 0.03) |
| Heavy atoms (residues 2-63) | 0.74 ± 0.18 (0.72 ± 0.09) |
| **RMSD from experimental restraints** | |
| NOE-based distance restraints (Å) | 0.0307 ± 0.0001 |
| Dihedral angle restraints (º) | 1.2 ± 0.02 |
| **RMSD from idealized covalent geometry** | |
| Bonds | 0.0063 ± 0.00002 |
| Angles | 0.671 ± 0.003 |
| Improper angles | 0.634 ± 0.005 |
| **Ramachandran analysis, residues 2-63 (%)** | |
| Residues in most favored regions | 93.1 |
| Residues in additional allowed regions | 6.9 |
| Residues in generously allowed regions | 0.0 |
| Residues in disallowed regions | 0.0 |
| *values in parentheses are mean global RMSD values (MOLMOL) | |

### <u>Supplementary Methods</u>

**Fragment analysis**

To evaluate agreement between sequence and structure for a given designed protein, we used Rosetta's standard fragment generation protocol[1] to select 200 fragments from natural protein crystal structures for each 9-residue-long segment of the designed protein. The fragments were chosen so that their sequence and secondary structure were as similar as possible to the sequence and *predicted* secondary structure of the designed protein segment (predicted using PSIPRED). Geometric similarity was quantified as the average RMSD of all 200 fragments at all positions (the "avg_all_frags" metric described further below*: Definition of scoring metrics*). Other measures of agreement are also described in that section.

**Adjustment of trypsin and chymotrypsin based on the "stability ladder"**

Using the "stability ladder" of previously measured stability scores for our 5 proteins, we adjusted the chymotrypsin values by a factor 3.5 to reproduce the previously reported stability data. A linear relationship was assumed. The stability cutoff was determined by plotting and fitting stability scores of the 2,300 random sequences.

**Definition of scoring metrics**

Simple sequence and topological properties:

description: the design name
sequence: the design sequence
dssp: the design secondary structure, according to the DSSP algorithm
n_res: the number of residues in the design
nres_helix: the number of helical residues in the design, according to DSSP
nres_sheet: the number of beta strand residues in the design, according to DSSP
nres_loop: the number of loop residues in the design, according to DSSP
frac_helix: nres_helix / n_res
frac_sheet: nres_sheet / n_res
frac_loop: nres_loop / n_res
n_charged: the count of D, E, K, and R residues in the designed sequence, plus one-half the number of H residues.
netcharge: the net charge on the design, assuming a charge of +1 on R and K, +0.5 on H, and -1 on D and E.
AlaCount: the count of Ala residues in the design
n_hydrophobic: the count of A, F, I, L, M, V, W, and Y residues in the design
n_hydrophobic_noA: the count of F, I, L, M, V, W, and Y residues in the design

Rosetta energy terms:

fa_atr, fa_dun_dev, fa_dun_rot, fa_dun_semi, fa_elec, fa_intra_atr_xover4, fa_intra_elec, fa_intra_rep, fa_intra_sol_xover4, fa_intra_rep_xover4, fa_intra_sol_xover4, fa_rep, fa_sol, hbond_bb_sc, hbond_lr_bb, hbond_sc, hbond_sr_bb, lk_ball, lk_ball_bridge, lk_ball_bridge_uncpl, lk_ball_iso, omega, p_aa_pp, pro_close, rama_prepro, ref, ss_sc, total_score, yhh_planarity: all the scores in the Rosetta full-atom energy function

Simple combinations of Rosetta energy terms:

score_per_res: total_score / n_res

fa_atr_per_res: fa_atr / n_res
fa_rep_per_res: fa_rep / n_res
hbond_lr_bb_per_res: hbond_lr_bb / n_res
hbond_lr_bb_per_sheet: hbond_lr_bb / nres_sheet
hbond_sr_bb_per_helix: hbond_sr_bb / nres_helix
net_atr_per_res: (fa_atr + fa_rep) / n_res
net_sol_per_res: (fa_sol + fa_elec) / n_res
net_atr_net_sol_per_res: net_atr_per_res + net_sol_per_res

Rosetta filters:
See
[https://www.rosettacommons.org/docs/latest/scripting_documentation/RosettaScripts/Filters/Filters-RosettaScripts](https://www.rosettacommons.org/docs/latest/scripting_documentation/RosettaScripts/Filters/Filters-RosettaScripts) for all documentation.
cavity_volume: void volume inside the designed structure, in $Å^3$, computed with CavityVolume filter
degree: average number of residues in a 9.5 Å sphere around each residue, computed with AverageDegree filter
contact_all: number of sidechain carbon-carbon contacts in the designed structure, computed with AtomicContactCount filter
exposed_hydrophobics: exposed nonpolar surface area of the designed structure, in $Å^2$, computed using TotalSasa filter, set to compute hydrophobic-only SASA
exposed_polars: exposed polar surface area of the designed structure, in $Å^2$, computed using TotalSasa filter, set to compute polar-only SASA
exposed_total: total exposed surface area of the designed structure, in $Å^2$, computed using TotalSasa filter
fxn_exposed_is_np: exposed_hydrophobics / exposed_total
holes: a normalized measure of the void volume inside the designed structure, computed with Holes filter
helix_sc: the average shape complementarity of each helical secondary structure element with the rest of the structure, computed using SSShapeComplementarity filter, set to evaluate helices only
loop_sc: the average shape complementarity of each loop element with the rest of the structure, computed using SSShapeComplementarity filter, set to evaluate loops only
mismatch_probability: the geometric average probability (across all positions in the design) that the designed residues will *not* adopt their designed secondary structures, as calculated by the PSIPRED algorithm from the designed sequence. Computed using the SSPrediction filter.
pack: a normalized measure of packing density, computed using PackStat filter
unsat_hbond: number of buried, unsatisfied hydrogen bonding atoms, computing using
ss_sc: the average shape complementarity of each helical or loop element with the rest of the structure, computed using SSShapeComplementarity filter
BuriedUnsatHbonds filter
unsat_hbond2: number of buried, unsatisfied hydrogen bonding atoms, computing BuriedUnsatHbonds2 filter

Custom metrics computed in Rosetta:
These metrics are not built-in Rosetta filters, but are computed within the Rosetta software

buried_np: buried nonpolar surface area in the designed structure on all amino acids, computed using version1 definitions of total nonpolar surface area per residue
buried_np_AFILMVWY: buried nonpolar surface area in the designed structure on nonpolar amino acids (AFILMVWY), computed using version2 definitions of total nonpolar surface area per residue
buried_np_AFILMVWY_per_res: buried_np_AFILMVWY / n_res
buried_np_per_res: buried_np / n_res
buried_minus_exposed: buried_np - exposed_hydrophobics
buried_over_exposed: buried_np / exposed_hydrophobics
exposed_np_AFILMVWY: exposed nonpolar surface area in the designed structure on nonpolar amino acids (AFILMVWY)
one_core_each: the fraction of secondary structure elements (helices and strands) with one large hydrophobic residue (FILMVYW) at a position in the core layer of the designed structure
two_core_each: the fraction of secondary structure elements (helices and strands) with two large hydrophobic residues (FILMVYW) at positions in the core layer of the designed structure
ss_contributes_core: the fraction of secondary structure elements (helices and strands) with one large hydrophobic residue (FILMVYW) at a position in either the core or interface layer of the designed structure
res_count_core_SASA: the number of residues in the core layer of the designed structure, with layers defined using solvent accessible surface area-based criteria
res_count_core_SCN: the number of residues in the core layer of the designed structure, with layers defined using sidechain neighbors-based criteria
percent_core_SASA: res_count_core_SASA / n_res
percent_core_SCN: res_count_core_SCN / n_res

<u>Custom metrics computed using external scripts as described in Rocklin et al. :</u>
abego_res_profile: Each position *i* in the designed structure can be classified by its ABEGO type, and the ABEGO types of positions *i*-1, *i*, and *i*+1 form a triad that defines the three-residue local structure at a coarse level. The abego_res_profile metric is the sum over all positions *i* in the designed structure of log ((p_aa | abego triad) / (p_aa)), where (p_aa | abego triad) is the frequency of the designed amino acid (from position *i*) in regions of natural proteins sharing the same ABEGO triad as the designed region centered on position *i*, and p_aa is the overall frequency of the designed amino acid at position *i*. At each position, this score is positive when the designed amino acid is overrepresented (compared with its normal frequency) in regions of natural proteins with the same local ABEGO triad structure as the designed region, and the score is negative when the designed amino acid is underrepresented in regions of natural proteins with the same local ABEGO triad structure.
abego_res_profile_penalty: Same as abego_res_profile, except summing over only positions with negative abego_res_profile scores (positions where the designed residue is typically underrepresented in the local structure).
contig_not_hp_avg: average size of the contiguous (in primary sequence) regions of the designed sequence lacking a large hydrophobic residue (FILMVWY)
contig_not_hp_norm: contig_not_hp_avg / (n_res / (1 + n_hydrophobic_noA))
contig_not_hp_max: the size of the largest contiguous region (in primary sequence) in the designed sequence containing no large hydrophobic residues (FILMVWY)

contig_not_hp_internal_max: the size of the largest contiguous region (in primary sequence) in the designed sequence containing no large hydrophobic residues (FILMVWY), excluding the regions between the first and last large hydrophobic residues and the termini
hphob_sc_contacts: the total number of sidechain-sidechain contacts between large hydrophobic residues (FILMVWY) in the designed structure
hphob_sc_degree: hphob_sc_contacts / n_hydrophobic_noA
largest_hphob_cluster: the size of the largest group of large hydrophobic residues (FILMVWY) that are all connected by at least one contact to each other in the designed structure
n_hphob_clusters: the number of disconnected groups of large hydrophobic residues (FILMVWY), where a group is defined as residues that contact each other in the designed structure but do not contact residues outside of the group
hydrophobicity: total hydrophobicity of the designed sequence, using the amino acid hydrophobicity scale from.

The column headers are annotated below in *Definition of scoring metrics*.

Fragment quality analysis:
Fragments were chosen for each designed protein using the standard Rosetta fragment generation protocol, which uses the designed sequence and PSIPRED-predicted secondary structure[2] as input. These metrics quantify the geometric agreement between the selected 9-mer fragments and the corresponding 9-mer segments of the designs (200 9-mer fragments are chosen per designed segment).
avg_all_frags: the average RMSD of all selected fragments to their corresponding segments of the designs, in Å. (200 × (n - 8) fragments in total)
avg_best_frags: the average RMSD of the lowest-RMSD fragment for each designed segment, in Å. (n - 8 fragments in total)
sum_best_frags: the sum of the RMSDs of the lowest-RMSD fragment for each designed segment. (n - 8 fragments in total)
worstfrag: among the set of fragments that are the lowest-RMSD fragments for their positions, the highest RMSD found
worst6frags: among the set of fragments that are the lowest-RMSD fragments for their positions, the sum of the RMSDs of the six highest RMSD fragments

**Feature expansion and RandomForest model**
Designs were analyzed using previous metrices and new features that combine connectivity and energetic terms which can be found in the score files as well as the jupyter notebook for predictions. Additionally, to previously reported score terms[3] we integrated the following new terms:
most_conRE: takes the Rosetta residue energy of the most connected residues as measured by how many amino acids are within 6 Å of the most connected residue.
most_conREn: takes the Rosetta residue energy of the most connected residues as measured by how many amino acids are within 6 Å of the most connected residue and then normalizes based on the number of neighbors
graph_density: measured the graph density of the contacting residues within the given protein.
bad_hub_penalty: extra penalty if well connected residue does not score well
highly_conREsum: total energy sum of the 4 most connected residues.

highly_conREsum_pres: highly_conREsum normalized by number of total residues in protein
highly_conREsumNorm: sum of residues energies of the top connected residues after normalization to total score of the protein
highly_conREsumNorm_pres: highly_conREsumNorm divided by total residues.
avgE_top_conResidues: average energy to top connected residues
avgE_top_conResiduesNorm: avgE_top_conResidues / total residues
avg_con: average connectivity within 6 Å
median_con: median connectivity
max_con: highest number of neighbors within 6 Å
num_bad_res: number of low scoring residues within the top connected residues.

**NMR spectroscopy and solution structure determination**

The main chain, and some side chain, chemical shifts were assigned using established triple resonance approaches employing a standard suite of experiments (($^{1}$H, $^{15}$N)-HSQC, ($^{1}$H, $^{13}$C)-HSQC, HNCA, HN(CO)CA, HNCACB, CBCA(CO)NH, HNCO, HN(CA)CO, HBHA(CBCACO)NH). Remaining side chain resonances were assigned using TOCSY, NOE, and aromatic-specific experiments (HCCH-TOCSY, H(CCO)NH-TOCSY, C(CO)NH-TOCSY, (HB)CB(CGCD)HD, $^{1}$H-TOCSY relayed constant-time ($^{1}$H, $^{13}$C) HMQC (aromatics), NOESY-($^{1}$H, $^{15}$N)-HSQC, NOESY-($^{1}$H, $^{13}$C)-HSQC, NOESY ($^{1}$H, $^{13}$C)-HSQC (aromatics)).

Manually identified peaks/signals in the 3D NOESY spectra were assigned and calibrated, distance restraints defined, and initial structures determined, in an iterative manner, using CYANA 2.1[4]. Using TALOS-N[5] main chain dihedral angle restraints and secondary structure predictions were derived based on assigned chemical shifts. The secondary structure predictions combined with results of initial structure calculations were used to define a set of hydrogen bond restraints for regions of well-defined secondary structure. These structural restraints were used to calculate a set of 500 initial structures starting from an extended structure using simulated annealing protocols in CNS 1.3[6] software suite for macromolecular structure determination[6, 7]. These were then refined and the 20 structures with the lowest energies selected as the final structural ensemble. CNS, MOLMOL[8], PyMOL, PROCHECK_NMR[9], and PROMOTIF[10] were used to analyze the ensemble and generate molecular models. Chemical shifts, structural restraints, and atomic coordinates have been deposited in the BMRB (entry 30844) and the PDB (PDB ID 7LDF).

**Thio-802 NMR structure analysis**

The amino acid sequence of the Thio-802 design used for structure determination by NMR spectroscopy included an N-terminal methionine residue and a C-terminal Leu-Glu linker followed by a six residue histidine affinity tag. The Leu-Glu linker and the first histidine residue could be observed by NMR, so the sequence used for structure calculations was 66 residues (N-terminal Met, 62 residues of the designed protein, C-terminal Leu-Glu-His). So, with regard to residue numbering, residues 1-62 of the designed protein correspond to residues 2-63 of the construct used for NMR.

Following refinement, the 20 Thio-802 structures, determined using NMR spectroscopy, with the lowest overall energies were chosen for final analysis. A summary of restraint information for the structure calculations and measures of overall structural quality for this 20-member ensemble are presented in Table S1. The overall RMSD for the main chain atoms of the ensemble is very low (0.1 Å, residues 2-63) indicating very good agreement for the atomic coordinates of the main chain for the members of the ensemble. The heavy atom RMSD is also very good (0.74 Å). There are no significant experimental restraint violations and more than 93% of the main chain dihedral angles are in the most favored regions of the Ramachandran space. These are all reliable indicators of high-quality structures.

As indicated in Figure S1, the most significant structural variability among the members of the Thio-802 NMR ensemble is in the loop between beta strands two and three (nominally residues P33-T36). For residues in this region, the relatively lower values for the Random Coil Index (RCI), a proxy for the model-free order parameter $S^2$, indicate increased motional mobility[11]. This appears to contribute to the lower restraint density observed for these residues and larger distance displacements from the mean (Fig. S1). These indicators suggest that this is a mobile, flexible loop with limited conformational restriction. The remaining loops/turns in the protein, and the C-terminal end of helix two, display similar, but far less pronounced, characteristics. The residues comprising the beta strands and alpha helices of the structures in the ensemble are generally well-restrained, display limited distance displacements from the mean, and display high RCI values, consistent with stable, well-ordered secondary structure of limited mobility.

A stereo view of the superposition of the main chain ribbon of the 20 members of the structural ensemble is shown in Figure S2. For most regions of the structure, the near perfect superposition of the main chains reflects the low RMSD and low distance displacements (Table S1, Fig. S1). The exception is the significant difference among the structures in the loop between beta strands two and three. A topology diagram based on the output of the HERA program[12] and various views of the ribbon diagram of the lowest energy structure of the NMR ensemble are also shown.
